# Neural signatures of natural behavior in socializing macaques

**DOI:** 10.1101/2023.07.05.547833

**Authors:** Camille Testard, Sébastien Tremblay, Felipe Parodi, Ron W. DiTullio, Arianna Acevedo-Ithier, Kristin L. Gardiner, Konrad Kording, Michael L. Platt

## Abstract

Our understanding of the neurobiology of primate behavior largely derives from artificial tasks in highly-controlled laboratory settings, overlooking most natural behaviors primate brains evolved to produce^1–3^. In particular, how primates navigate the multidimensional social relationships that structure daily life^4^ and shape survival and reproductive success^5^ remains largely unexplored at the single neuron level. Here, we combine ethological analysis with new wireless recording technologies to uncover neural signatures of natural behavior in unrestrained, socially interacting pairs of rhesus macaques. Single neuron and population activity in prefrontal and temporal cortex unveiled robust encoding of 24 species-typical behaviors, which was strongly modulated by the presence and identity of surrounding monkeys. Male-female partners demonstrated near-perfect reciprocity in grooming, a key behavioral mechanism supporting friendships and alliances^6^, and neural activity maintained a running account of these social investments. When confronted with an aggressive intruder, behavioral and neural population responses reflected empathy and were buffered by the presence of a partner. By employing an ethological approach to the study of primate neurobiology, we reveal a highly-distributed neurophysiological ledger of social dynamics, a potential computational foundation supporting communal life in primate societies, including our own.

## MAIN TEXT

Survival and reproductive success for long-lived social primates like humans, apes and monkeys, depends upon deft deployment of cooperation and competition, often by forming friendships and alliances that provide support in the face of social and environmental challenges^5,7,8^. In humans, social deficits severely impact daily life function for individuals with disorders like autism and schizophrenia, and the inability to connect with others is associated with depression and other long-term negative impacts on health and well-being^9–11^. Despite their importance, the neural circuit mechanisms supporting investments in long-term social relationships remain poorly understood.

For practical reasons, investigations of the neural basis of human and macaque social behavior have been restricted to highly-controlled laboratory settings in which individuals interact socially via an interposed computer, often in the context of economic games^12–19^. Despite the utility of this approach, which provides the opportunity to link neural signals to mathematically tractable variables, it neglects the natural behaviors primate brains evolved to produce and the social contexts that determine their function^2,3^.

In this study, we focus on the neurobiology of dynamic social interactions in rhesus macaques (*Macaca mulatta*) whose behavior and biology has been better characterized than any other primate species^20^. Moment-to-moment decisions to travel, forage, fight, groom, mate or rest in rhesus are strongly influenced by the presence, identity, and actions of other conspecifics^4,21^. Macaques form friendships and alliances, in part built and maintained through reciprocal grooming, enabling mutual support to gain access to limited resources or fend off threats^6^. These observations imply that macaque brains encode rich information about the social environment and integrate this information over multiple timescales to guide adaptive decision making.

Prior laboratory studies suggest social information processing in the primate brain proceeds hierarchically, from the perception of visual and auditory social cues in temporal cortex^22,23^, to integration of others’ actions, intentions and rewards in the orbitofrontal, cingulate, and prefrontal cortex^14,17,18,24^. Here, we endeavored to identify how this system encodes information during unconstrained interactions between macaques. We focused on two areas located at two ends of the social information-processing hierarchy: visual processing area TEO^22,25,26^ and socio-cognitive processing area 45 in the ventrolateral prefrontal cortex (vlPFC)^24,27,28^. These areas were chosen both for their established roles in sensory and cognitive processing of social information, respectively, and because they are on the gyral surface and therefore accessible to our novel wireless recording technology supported by Utah arrays.

We predicted firing patterns of vlPFC neurons would contain rich information about individual behaviors and the social environment robust to visuo-motor contingencies. By contrast, we predicted TEO neurons would be highly sensitive to changes in visual orientation, and would not reflect current social and behavioral contexts. Neuronal activity in these areas has never been studied in unrestrained primates engaging in natural social interactions before, so the extent and nature of information encoding remained open and not readily predictable based on prior laboratory work. For example, given the importance of reciprocal investments in relationships through mutual grooming for rhesus and other primates, we postulated that vlPFC might maintain a running account of these interactions–a prediction never tested before using standard laboratory task-based paradigms.

Here, we used neural data-logging technologies to record from hundreds of neurons in the cortex of freely-moving, socially-interacting pairs of rhesus macaques. During hours-long sessions, we paired male subject monkeys with their female partners or isolated them, manipulated the identity of neighboring monkeys, introduced social stressors, and provided opportunities for expression of macaques’ natural behavioral repertoire, including non-social behaviors like foraging, locomotion, and rest. Combining ethological, computer-vision and neural population analytical methods, we discovered widespread, robust neural signatures of 24 natural behaviors and their social context, in both areas, which were not reducible to sensorimotor contingencies. Remarkably, neurons in both brain areas tracked dynamic investments in grooming given and received during mutual interactions. Moreover, neural responses reflected social support and empathy in response to an external stressor. Together, our findings reveal a highly-distributed neurophysiological ledger of social dynamics unfolding across multiple contexts and extended timescales, a potential neural foundation supporting the affiliative bonds that drive survival and reproductive success in gregarious primates^7^, including humans ^9^.

### Unrestrained macaques recapitulated their natural behavioral repertoire

Subjects were two male rhesus macaques interacting spontaneously with their respective female partners over 2.5-hour long sessions in their home enclosure (N = 12 sessions, 6 per subject; **Fig 1A**). In each session, we provided monkeys with foods requiring manual extraction from a turf board (i.e. foraging) and experimentally manipulated social context pseudo-randomly in a block design by: 1) isolating or pairing the subject with his female partner, and 2) when paired, varying the identity of neighboring monkeys (total: 3 conditions; **Fig 1A**). Neighbors were monkeys in the adjacent enclosure that were visible but not physically accessible. The behaviors we focused on occurred over seconds to minutes and were labeled manually from videos captured by 7 cameras, at 1 second resolution, by two primatologists with extensive field experience (93% inter-rater reliability, see **Methods**) using a well-validated ethogram^8^ derived from observations of free-ranging rhesus macaques (**Appendix 1-2**).

**Fig 1.**
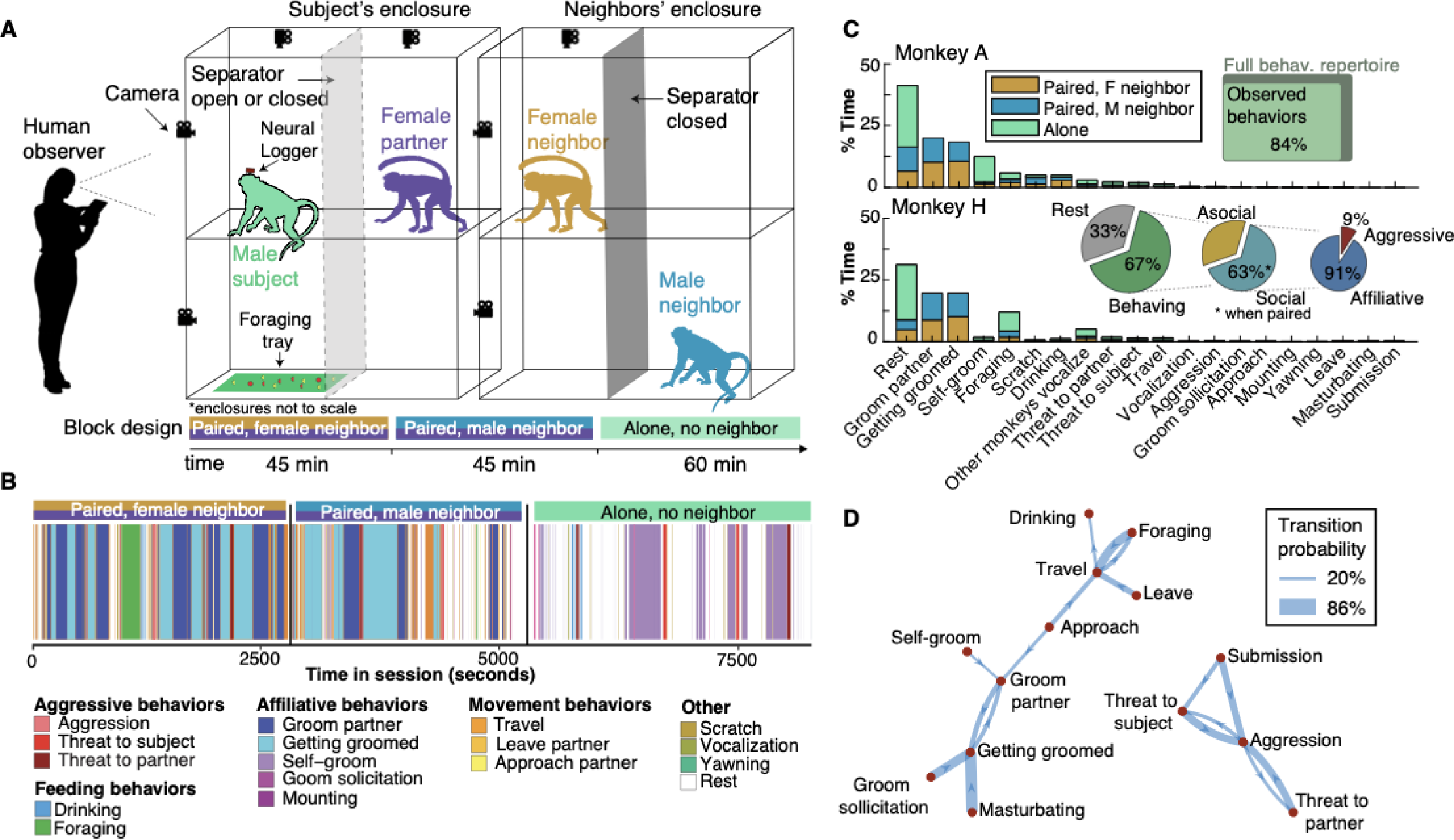
Unrestrained socially-interacting macaques spontaneously express their species-typical behavioral repertoire. (**A**) Experimental setup with two male-female pairs in adjacent enclosures. Neural activity was recorded from 2 male subjects during various social contexts and behaviors in 2.5-hour sessions (N=6 sessions per subject). (**B**) Species-typical behaviors expressed during 3 social contexts during a typical recording session (N=24 behaviors over 12 sessions). (**C**) Proportion of time engaged in each behavior across all 12 sessions, for each male monkey. When paired, monkeys engaged in social behaviors with their partner 63% of the time. (**D**) Behavioral state transition probabilities (thresholded >20%) segregate into affiliative (left) and aggressive (right) clusters (see **Fig S1B** for full behavioral transition matrix). Monkey and human silhouettes in (A) created with BioRender.

Monkeys displayed nearly the full repertoire of aggressive, affiliative, locomotive, and feeding behaviors observed in the wild (N=24 behaviors, 84% overlap with wild macaque ethogram, **Fig 1B-C**). While paired, monkeys spent most of their time grooming (>60%, **Fig 1C**), defined as repetitive brushing through another monkey’s fur. For macaques, like other primates, grooming serves as an investment that builds and maintains strong social bonds^29^. Male-female pairs engaged in reciprocal grooming interactions **(Fig 1B**, alternating light blue and dark blue). Aggression constituted a small but significant portion of social interactions (9%) and was most frequently expressed in response to threats from human intruders. Rest, defined as sitting and observing the environment passively without engaging in any other overt actions, occurred 33% of the time.

Transitions from one behavior to another were highly-structured, with some behaviors followed by specific behaviors with high probability (**Fig 1D, S1A-B**). For example, a monkey soliciting another monkey to groom himself elicited a grooming bout with 68% probability, whereas a subject monkey being threatened provoked an aggressive response with 60% probability. Behavioral transitions coalesced into affiliative (left) and aggressive clusters (right; **Fig 1D**), establishing two major affective contexts for behavior.

### Single neuron activity is broadly modulated by species-typical behaviors

We developed and deployed new technology that interfaced chronically-implanted, multi-electrode Utah arrays (Blackrock Neurotech, UT) with miniaturized, wireless data-loggers (Deuteron Technologies, Israel) to record from approximately 300 units (single and multi-units) per session (**Fig 2A**, mean = 307.42 units, SD = 19.67) while monkeys moved and interacted freely in their home environments (**Movie S1-4**). This technology allowed us to obtain stable, high-quality neural recordings unencumbered by wires or electromagnetic noise, while minimizing the cranial implant footprint (**Fig S1C-D**, see **Methods** for more details). We analyzed firing rates of individual neural units in 1 sec bins to match the timescale of behavioral quantification by observer primatologists (see **Methods**).

**Fig 2.**
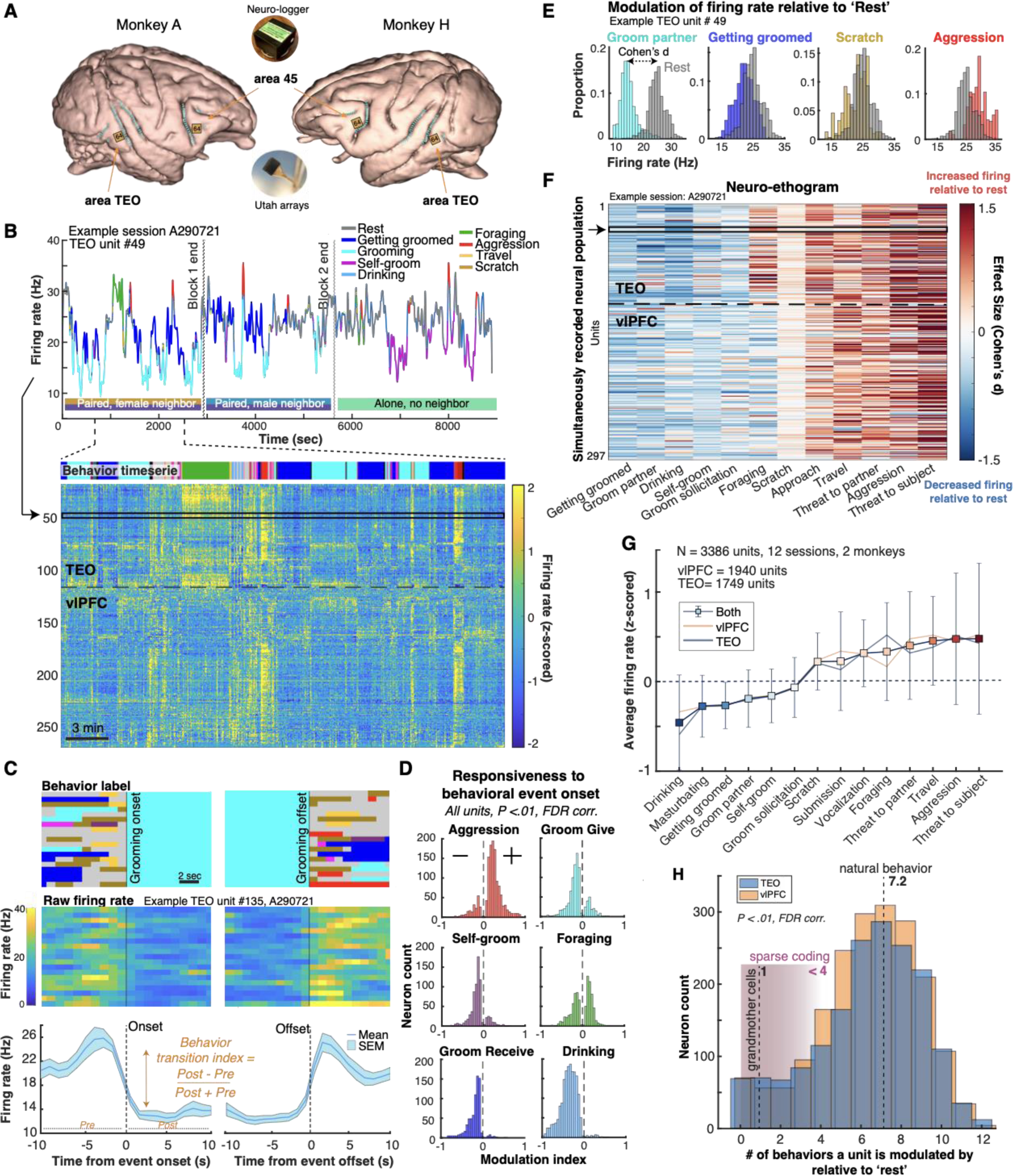
Single unit activity varies across the natural behavioral repertoire. (**A**) 3D MRI reconstruction of subjects’ brains and recording locations. Two 64-channel Utah arrays were implanted in the right hemisphere for monkey A and left hemisphere for monkey H. Data was stored on a wearable microSD card using a data-logger (inset). (**B**) Top panel: Example TEO unit firing rate trace during an example 2.5-hour session with the subject’s behavior labeled. Bottom panel: Z-scored firing rate over the entire population of simultaneously recorded units for ∼33min of the session (See **Fig S2A** for full session), y-axis ordered with hierarchical clustering using rastermap^37^. (**C**) Top: Behavioral labels of all instances of grooming partner in an example session, from 10 sec before to 10 sec after grooming onset (left) and offset (right). Middle: Raw firing rate for an example neuron aligned to grooming onsets and offsets. Bottom: Average firing rate across all grooming instances for the same neuron as in the middle panel. (**D**) Distributions of behavior transition indices for units showing significant modulations in firing rate by behavior (*t*-test pre vs. post-activity, FDR-corrected, *P* < 0.01), for activity aligned to the onsets of aggression, grooming, foraging, self-groom, getting groomed and drinking. (**E**) Normalized histograms of the distribution of firing rates during behaviors compared to ‘rest’ (reference behavior; gray) for example unit in panel B. (**F**) Neuro-ethogram plot displaying effect sizes (Cohen’s d) of difference in distributions of firing rate during each behavior (columns) which occurred for at least 30 sec in the session compared to rest (as in C), for all units (rows), during an example session. Example unit from panels B and C shown (black arrow). Red: excitation; Blue: suppression. **G**) Averaged z-scored firing rates of all TEO and vlPFC units by behavior, pooled across monkeys. Behaviors ordered from low to high average firing rate. Error bars are SD. (**H**) Distribution of number of behaviors for which each neuron was modulated by compared to rest (*t*-test, FDR-corrected, *P* < 0.01), for TEO and vlPFC, pooled across animals. Sparse coding (pink shading) predicts each neuron to encode < 4 behaviors (**Table S1**), while “Grandmother cell hypothesis” predicts selectivity for a single behavior.

Over the course of a session, neural activity in both TEO and vlPFC showed striking, continuous modulation by behavioral state (**Fig 2B, S2A,F**), with responses aligned to behavioral transitions (**Fig 2C**). We quantified individual units’ responsiveness to behavioral events by computing a “behavior transition index”, comparing activity before and after behavior onsets (**Fig 2C,D**). Behavior transition indices range from -1, corresponding to strong activity suppression following the onset of a behavior, to 1, corresponding to strong activation. Because this study did not follow a standard trial-based design, repeated sequences of behaviors within a session were relatively rare (**Fig S1A**), such that we used a complementary approach to quantify unit responsiveness. We quantified deviations in neural activity from a reference state of “rest” using an effect size metric (Cohen’s d) comparing firing rate distributions of behaviors to rest (**Fig 2E**). In a “Neuro-ethogram” plot, we depict differences in mean firing rates between behavioral states and rest (i.e., effect size) for an example sample of simultaneously-recorded neurons (**Fig 2F**). Units tended to collectively decrease firing rates during affiliative behaviors and increase firing rates during aggression-related behaviors and travel relative to rest (**Fig 2G**). Surprisingly, patterns of neuronal modulation in TEO and vlPFC were highly similar. A noteworthy exception is modulation during foraging in the example session shown in **Fig 2B**. This between-area difference in activity was only observed in a subset of sessions in one monkey (see **Fig S2F** for more sessions which show sub-populations of neurons differentially modulated during foraging in both areas).

In both brain areas, firing rates of most neurons were significantly modulated by a large array of natural behaviors (median = 7.2 behaviors, **Fig 2H**; see statistically thresholded neuro-ethogram in **Fig S2B**). This finding held even when restricting analyses to well-isolated single neurons and controlling for sample size (**Fig S2C-D**). Although mixed selectivity for 2-3 task variables has been reported in many laboratory tasks^30–32^, the multiplexing we observed during natural behavior exceeds reports from prior controlled laboratory experiments^33,34^ (**Table S1**). Together, these findings support distributed rather than sparse coding models of cortical activity in primates, inviting reconsideration of theories that propose sparse coding as a mechanism that balances metabolic constraints with information-processing capacity^35,36^.

### Neural population activity clusters along behaviors and social contexts

We next examined activity across the population of simultaneously recorded neurons, which can reveal information not detectable in the average activity of individual neurons^38,39^. We applied multivariate analytical methods to investigate neural signatures of the diversity of natural behaviors and their variation across social contexts in vlPFC and TEO population activity. First, we visualized neural population activity patterns using an unsupervised nonlinear dimensionality-reduction technique, *Uniform Manifold Approximation and Projection* (UMAP)^40^. Nearby points in this dimensionality-reduced space represent similar neural states and this visualization provides insights into how neural states are structured as a function of behavior and social context.

When plotting the first 3 UMAP axes, we observed that neural population activity in vlPFC spontaneously clustered along species-typical behavioral states such as grooming, foraging and aggression (**Fig 3A**, see **Fig S3A-B** for plots including all behaviors). Surprisingly, neural activity in TEO also spontaneously clustered along behavioral states (**Fig 3B**). Patterns that were not obvious from analyzing activity of individual units emerged clearly when considering population activity. For example, grooming and being groomed were both associated with suppression of mean firing rates of individual units relative to rest (see **Fig 2D**), but segregated into distinct clusters when the simultaneous activity of all recorded units was considered (**Fig 3A,B**).

**Fig 3.**
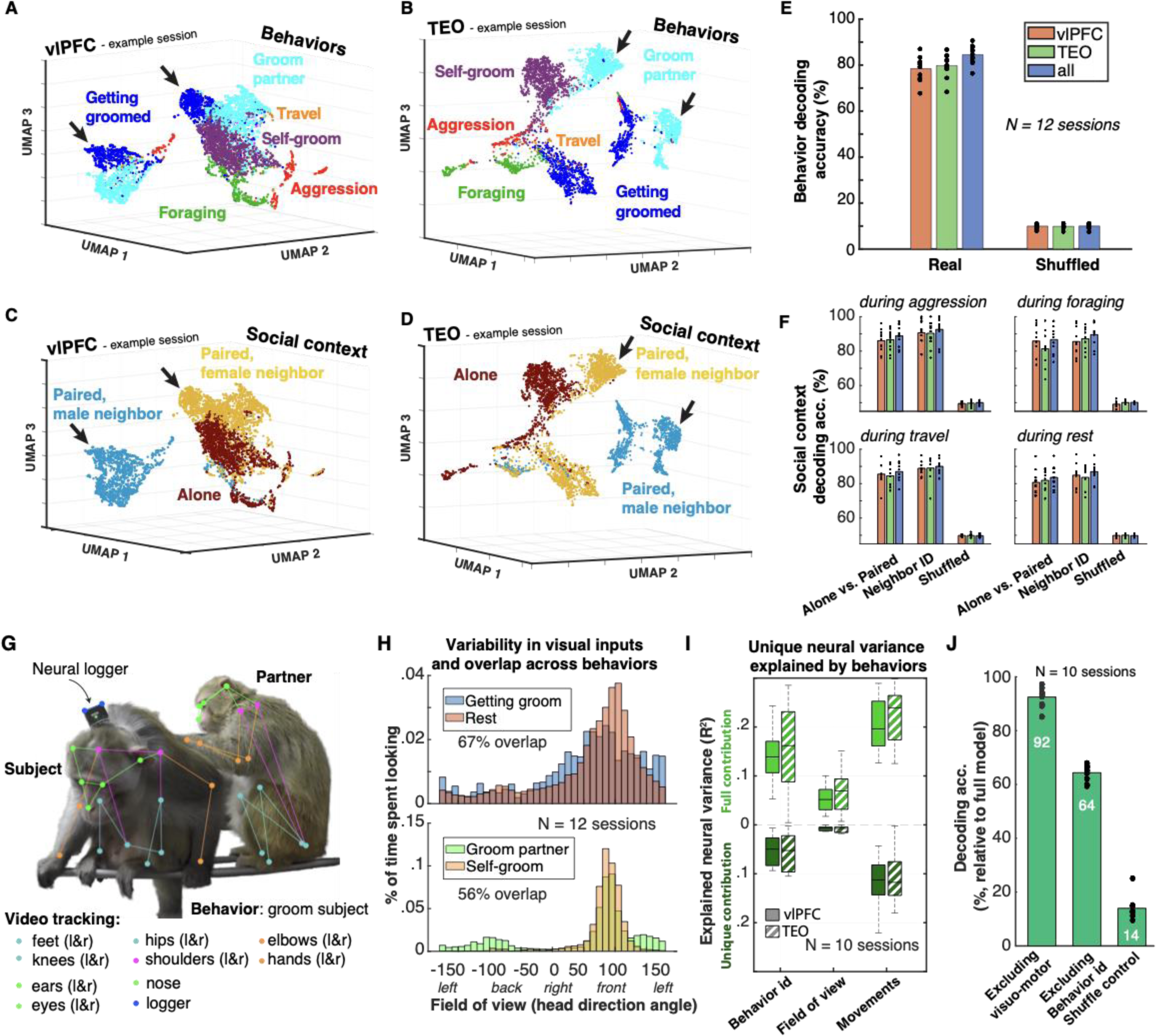
Neural population activity simultaneously encodes current behavior and social context. (**A**) Neural population states in one example session visualized using unsupervised UMAP, color-coded post-hoc by subject’s behavior. Each point represents simultaneous firing of all vlPFC (A) or TEO (B) units for a 1 second bin. For visualization, only behaviors that occurred for at least 30 sec within a session are plotted and “rest” is excluded (see **Fig S3 A-B** for a plot including all behaviors, **S3C-D** for Euclidean distances within and between behavior states & contexts in UMAP space across all sessions, and **S3E-H** for UMAP projections using different parameters). (**C-D**): Same UMAP space as in A-B but color-coded by social context (i.e., alone, paired & female neighbor, paired & male neighbor). Black arrows indicate example behavioral sub-clusters that segregate according to context. (**E**) Decoding accuracy for behaviors across sessions, only considering behaviors that occurred for at least 30 seconds in a session (mean N= 12 behaviors on any given session, chance = ∼8%). Each dot represents a session. (**F**) Decoding social context during aggression, foraging, travel and rest separately. Chance is ∼50%, same color-code as in E. (**G**) Schematic of movement quantification including 17 body landmarks, head position, and head direction (extracted from neural logger position). (**H**) Histograms of field of view during getting groomed vs. rest (top) and groom partner vs. self-groom (bottom) showing overlap. (**I**) Neural variance explained by behavioral states (Behavior ID), field of view, and movements obtained by regressing population activity onto those variables over the time-course of an entire session. (**J**) Decoding accuracy of an SVM decoder predicting the behavioral ethogram (as in **Fig 3E**) from neural population activity, while controlling for visual and motor confounds. Scores relative to the full model shown in **Fig 3E**. Note that we could not perform video tracking on 2/12 sessions due to missing videos from appropriate angles (see **Methods**). Thus, panels E-F contain data from 10 sessions.

Furthermore, we found that population state clusters for most behaviors were divided into sub-clusters. For example, the clusters ‘Groom partner’ and ‘Getting groomed’ (black arrows in **Fig 3A,B**) split into two sub-clusters, in both areas vlPFC and TEO. When we recolored each UMAP plot according to social context instead of behavior, we found these subclusters were explained by the social context in which those behaviors occurred (**Fig 3C,D**). For example, the cluster ‘Groom partner’ (black arrows in **Fig 3B**) split into two sub-clusters corresponding to whether the neighbor was a male or a female during the grooming bout (black arrows in **Fig 3C** indicate the same sub-clusters). Thus, the structure of neural population activity also reflected social context (see **Fig S3C-D** for replication across sessions). Simultaneous representation of both actions and contexts may be a mechanism by which neural activity tracks and guides context-dependent behavioral choices in socially-interacting primates^18,41^.

To complement UMAP visualizations of the data, we applied a widely used linear decoding technique (Support Vector Machine^42^, see **Methods**) to determine what information a hypothetical downstream brain region could decipher from neural activity in TEO and vlPFC. Natural behaviors were highly decodable on a moment-by-moment basis from simultaneously recorded neural activity of both brain areas, together and from each independently (mean decoding accuracy 84%, N=12 behaviors with >30 samples per session, **Fig 3E**). Social context was also linearly decodable (mean accuracy >80%, **Fig 3F**) from one or both brain areas during various behaviors. It was possible, for example, to decode whether the subject monkey was foraging alone or with his partner, or whether the neighboring monkey was a male or a female during a bout of aggression. Together, these findings resonate with decades of field observations of primates demonstrating the importance of social context, such as the identity of neighboring monkeys, in the decision-making processes that drive behaviors like socializing, foraging, or traveling^43,44^.

Movements dominate cortical dynamics in mice^45,46^ and explain a large portion of neural activity in macaque cortex during laboratory task performance^32^. Moreover, area TEO has long been linked to visual processing of objects and social stimuli^26^. These observations raise the question of whether the patterns of neural population activity we uncovered reflect movements or changes in visual inputs associated with specific behaviors and social contexts rather than the behaviors or contexts themselves. To address this possibility, we tracked movements and field of view from video using deep learning approaches (**Fig 3G**, see **Methods**) and quantified their relationship to neural activity using complementary analytical methods. First, we found that head direction (proxy for field of view) varied substantially within a given behavioral state and the same visual inputs and movements occurred across different behaviors and contexts, partially decorrelating behaviors from visuo-motor contingencies (**Fig 3H, S3K**). Despite visuo-motor similarities, neuronal activity was still distinctively modulated by behavioral states and social contexts (**Fig3A-F**). Second, using multi-linear ridge regression^32,45^, we quantified the relative neural variance explained by behavioral states, movements (17 key points across the whole body, **Fig 3G**), and head direction (see **Methods**). After accounting for movements and changes in field of view (proxied by head direction), behavioral states still uniquely explained part of the neural variance in both brain areas (**Fig 3I**). Finally, after regressing out visuo-motor variables from neural activity, decoding accuracy for behavior decreased by only 7.5% on average, whereas regressing out ethologically-defined behavioral states decreased accuracy by 35.6% (**Fig 3J**). Taken together, these findings indicate that patterns of neural activity in TEO and vlPFC at least partially represent species-typical behavioral states and social contexts independent of immediate visuo-motor contingencies. We also note the presence of strong movement signals in freely-moving primates, even in an extrastriate visual area like TEO, which has important implications for models of how brains function during natural behavior^47^.

### Neural population activity tracks grooming reciprocity

Strong, stable relationships in humans and other animals are maintained through reciprocal interactions^48–51^. In macaques and other primates, reciprocity takes the form of mutual grooming (**Fig 4A**) and support in the face of challenges^6^ and can occur over days to months^50,52^. It has been proposed that an internal “mental account” tracks these mutual social investments over long periods of time (“calculated reciprocity”^53^). We next investigated the behavioral and neural mechanisms supporting grooming reciprocity over time in our two male-female pairs.

**Fig 4.**
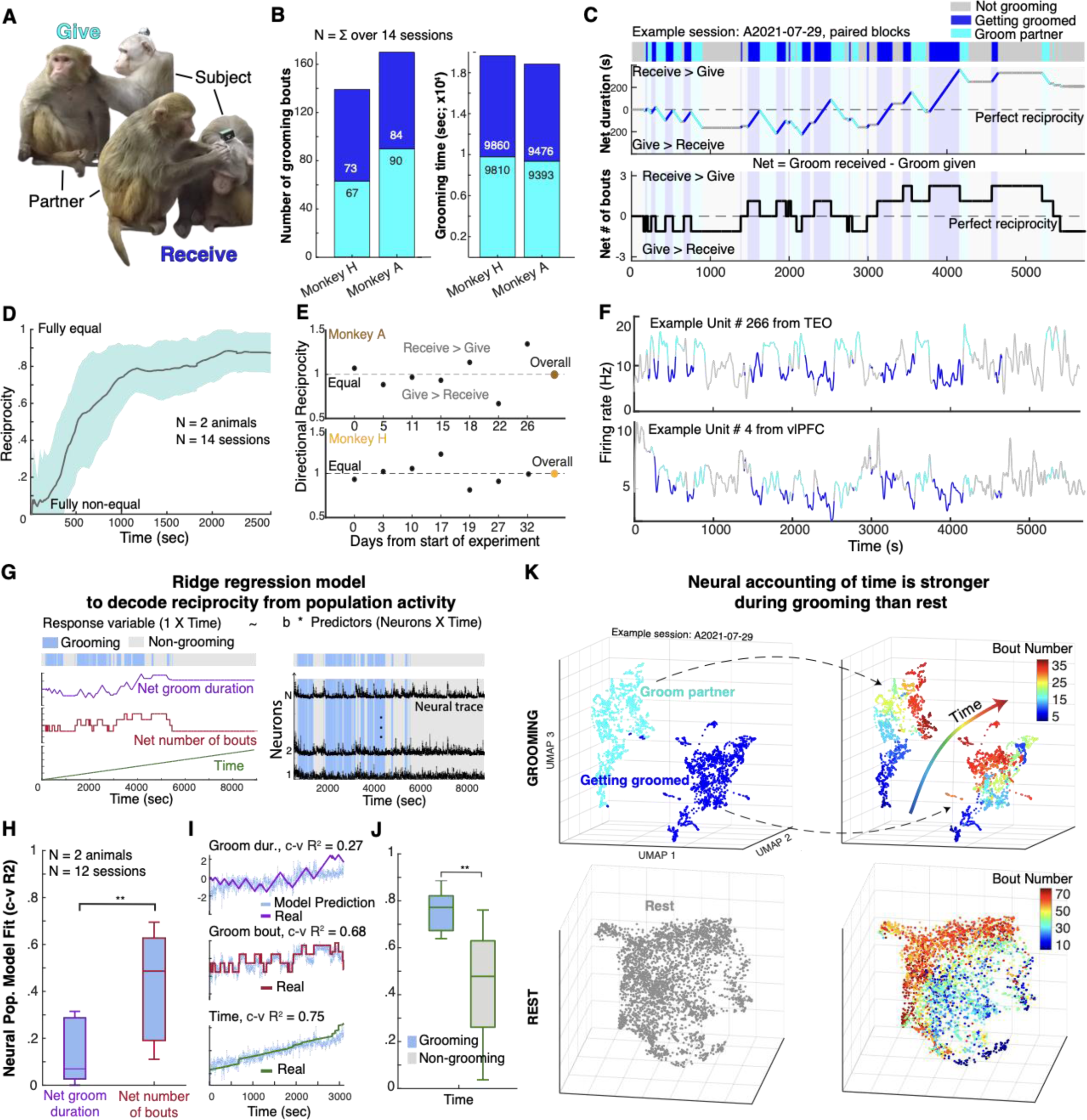
Neural population activity tracks grooming reciprocity. (**A**) Images of reciprocal grooming between subject and partner in one pair of monkeys. (**B**) Number of grooming bouts (left) and grooming time (right) given (cyan) and received (dark blue), for both subject pairs, summed across all sessions. (**C**) Reciprocal grooming for an example session. Top: grooming bouts over time, received (dark blue) or given (cyan). All non-grooming behaviors are lumped into a “not grooming” category (gray). Middle: Net grooming duration in seconds in the session. Bottom: Net number of grooming bouts in the session. Net = Received - Given. Dashed lines indicate perfect reciprocity. (**D**) Reciprocity in grooming duration over time during a session, averaged across sessions. Shaded area represents standard errors. (**E**) Directional reciprocity (y-axis) across sessions which were separated by multiple days (x-axis). Directional reciprocity goes from 0 to 2, with 1 being equal grooming duration, <1 more groom given than received, >1 more groom received than given (see **Methods**). (**F**) Raw activity of two example units from the same session as in panel C., groom receive (dark blue), groom give (cyan). (**G**) Schematic of ridge regression approach to predict net grooming duration, net number of bouts and time elapsed within a session from the neural population activity. (**H**) Cross-validated model fit (R^2^) for net duration and net number of bouts models using ridge regression. Predictions based on neural activity during grooming only. (**I**) Model predictions and real values for net grooming duration (top), net number of bouts (middle) and elapsed time (bottom) for same example session A. (**J**) Cross-validated model fit (R^2^) for predicting elapsed time during grooming and outside of grooming. (**K**) Unsupervised UMAP projection of neural population activity during grooming only (top) vs. rest only (bottom), pooled across both brain areas. Left: neural states in UMAP space color-coded by behavior. Right: color-coded by chronological bout number in this session. Bout numbering in grooming includes both directions (give and receive). Note that 14 sessions were used for panels A-E and 12 sessions for panels H & K. Note that 2 sessions without neural data were included in the behavioral analyses of panels A-E to include all consecutive grooming interactions in this experiment which was critical to unveil the temporal dynamics of reciprocity over multiple days (see **Methods**).

Combining grooming interactions across all consecutive sessions, both male-female pairs showed remarkable reciprocity in both the number of grooming bouts and total duration of grooming (**Fig 4B**). The observed level of reciprocity in bout quantity (>0.96 in both subjects, 1=perfect reciprocity and 0=fully one-sided, see **Methods**) was highly unlikely to occur by chance (*P* < 0.0001 for both pairs, **Fig S4A** top panel). Monkeys tended to take turns grooming (i.e., alternated in quick succession, *P* < 0.001 for both pairs, **Fig S4A** middle panel, **Fig S4C**), which can facilitate tracking the number of bouts to achieve reciprocity, but did so in less than 50% of grooming interactions. These findings suggest monkeys tracked the number of bouts given and received over multiple timescales to achieve observed levels of reciprocity.

Monkeys also tracked grooming duration. The observed near-perfect grooming duration reciprocity (>0.98) would be unlikely to occur by chance (*P* < 0.05 across both monkeys, **Fig S4A** bottom panel). One simple heuristic to achieve equal duration is to match the length of the immediately-preceding grooming bout. We did not find a correlation between the durations of consecutive alternating grooming bouts (r=0.04, p>0.7, **Fig S4B**), suggesting monkeys did not use this heuristic. Instead, reciprocity was progressively achieved over the course of the study, both within 2.5 hour-long sessions (**Fig 4C-D**) and across multiple days (**Fig 4E**). Remarkably, small grooming duration imbalances at the end of one session were rebalanced in subsequent sessions, which could occur up to 7 days later (**Fig 4E**). Taken together, these observations suggest monkeys maintain a running “mental account” of both the quantity and duration of grooming bouts given and received, over extended periods of time, to achieve reciprocity–even in the laboratory. This exceptional cognitive competence has not been previously documented in monkeys with the temporal resolution afforded by our study and, in this context, supports “calculated reciprocity”^53,54^.

We investigated potential neural signatures of this mental accounting of reciprocity and found single units in both TEO and vlPFC that modulated their firing rates according to whether the subject was giving or receiving grooming (**Fig 4F**). Next, we asked how population activity in TEO and vlPFC tracked net number of grooming bouts and grooming duration (Net = Receive - Give, **Fig 4C**), as well as elapsed time within a session as a potential confound since it was, in some sessions, correlated with net groom time (**Fig S4E** top panel). We used multi-linear ridge regression analysis with 5-fold cross-validation to predict these 3 continuous variables from neural population activity (**Fig 4**, see **Methods**). We found that population activity in both brain areas and both monkeys predicted net grooming duration and net number of bouts within a session (mean R^2^ net duration = 0.16, R^2^ net bout number = 0.47, **Fig 4H-I**, **S4F-G**). Model predictions for net grooming duration, however, were much weaker than for net number of bouts, and could be explained by elapsed time within a session (**Fig S4E**). Net grooming bout, by contrast, did not correlate with elapsed time and was also highly decodable from neural population activity (>80% accuracy, **Fig S4H**). We therefore hypothesized that long-term mental accounting for grooming reciprocity occurs by integrating elapsed time with net number of bouts given vs. received, rather than tracking net time spent grooming. No single unit on its own could predict these reciprocity metrics as well as the neural population (**Fig S4I**). Moreover, the distribution of single unit model fits for all three models (duration, bout and time, **Fig S4I**) were indistinguishable, even though they significantly differed when considering the neural population as a whole (**Fig 4H-I**, **Fig S4G)**. These findings suggest that a highly distributed neural code supports mental accounting of grooming bouts and elapsed time in order to achieve calculated reciprocity.

Notably, neural tracking of elapsed time was stronger when monkeys were grooming than when they were not (**Fig 4J**). UMAP visualizations of neural activity during grooming showed striking temporal structure defined by bout number, whereas UMAP visualizations of neural activity during rest were not temporally structured to the same extent (**Fig 4K**). This difference was also reflected in higher linear decoding performance for chronological bout number during grooming compared to rest (mean accuracy during grooming: 89%; rest: 59%; chance: 10%, **Fig S4F**). This result provokes the hypothesis that grooming, which is both rewarding and holds unique functional significance for building and maintaining relationships, enhances the salience and importance of elapsed time. Overall, our findings reveal a potential neural mechanism by which reciprocity is tracked in social relationships.

### Neural responses to threats reflect empathy and social support

Social relationships help primates cope with external stressors, such as environmental disruption^8^ or aggressive encounters with conspecifics^55^. The presence of a supporting partner (i.e., social support) is associated with lower physiological stress response to threats^56,57^. One important emotional driver of this social support is empathy– affective resonance with the feelings of a partner^58^. Here, we investigated the neural correlates of both social support and empathy using a standard “human intruder” paradigm designed to elicit aversive responses (**Fig 5A**). We analyzed behavioral and neural responses to 30 seconds of open-mouth threats (a particularly aversive stimulus for macaques^59^) directed towards the subject monkey or his partner, while the subject was either alone or paired.

**Figure 5.**
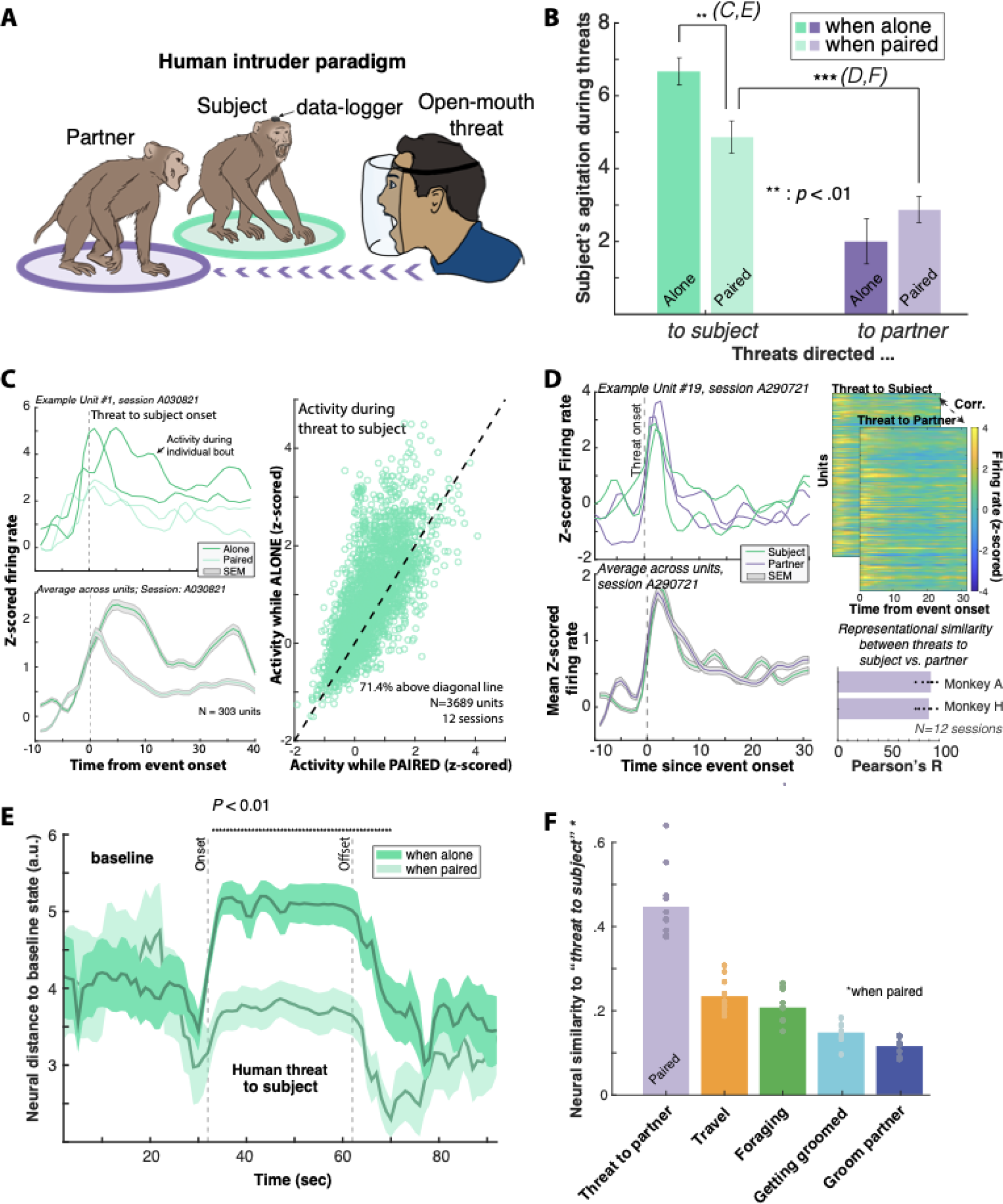
Behavioral and neural evidence of social support and empathy in response to threats. (**A**) Schematic of the “human intruder” paradigm (see **Methods**) where, in this example, the partner is being threatened by an experimenter with an open-mouth threat for 30 seconds. Both the partner and subject monkeys were threatened while alone, or with a partner (**B**) Behavioral responses to human threats towards the subject or the partner, when alone (dark bars) or paired (light bars). Agitation measured by manual scoring of videos (details in **Methods**). (**C**) Top left: neuronal activity of an example unit during threats to the subject when paired (light green) vs. alone (dark green). Bottom left: Average response to threats towards the subject across all recorded units for an example session. Same color code as in top left. Right: mean z-scored activity during threats towards the subject while paired (x-axis) vs. alone (y-axis). Each point represents one unit and we show data across all recorded units. 71.4% of units respond more to threats while alone. (**D**) Left panels: same as C but comparing threat toward the subject vs partner when paired. Top right: z-scored average neural population responses during threat to subject compared to threat to partner while paired, for an example session. Bottom right: Representational similarity scores, i.e. correlation coefficients between average neural population responses during threat to subject and threat to partner while paired. Each dot represents one session. (**E**) Mean Euclidean distance from baseline (center of mass of ‘rest’ states) in UMAP neural space during threats towards the subject in the alone (dark green) vs. paired conditions (light green), averaged across threat bouts (N = 25 alone, N = 24 with a partner). Shading represents SEM. (**F**) Similarity score in UMAP space between neural states during threat to subject and threat to partner (purple), and other reference behaviors. Artwork in (A) by Jordan Matelsky.

Open-mouth threats provoked aversive reactions by our subjects, including aggressive displays, attacks, and submissions, which we combined into an “agitation” score. We found that agitation was strongest when subjects were alone, and the presence of a partner reduced agitation by 27% (**Fig 5B**, two-sample t-test; *t*(1, 36) = 2.9, *P* = .006, see **Fig S5A** for results separated by monkey). Similarly, individual neural responses to threats were dampened when subjects were paired with a female partner (**Fig 5C**). At the population level, the Euclidean distance of threat states in UMAP neural space relative to ‘rest’ was higher when the subject was alone compared to when he was paired (**Fig 5E**, *P* < .01, see **Fig S5B** for results separated by monkey). Notably, female partners consistently groomed their male partner following threats, consistent with the known stress-reducing functions of this affiliative behavior (**Fig S5C**). When the subject and the partner were paired, threats directed towards the partner elicited defensive reactions from the subject, albeit to a lesser extent than threats directed to himself (**Fig 5B**). Threats towards the subject or his partner elicited remarkably similar patterns of neural activity, both at the single unit level (**Fig 5D** mean representational similarity across sessions *r* = 0.91 for monkey A and *r* = 0.90 for monkey H) and the population level (**Fig 5F**, threat states are closest in UMAP space compared to other behaviors). The similarity of neural responses could not be explained by common visual inputs; neural activity in both conditions was correlated whether the subject was looking towards or away from the threatening stimulus (**Fig S5D-E**). This neural “mirroring” between threats directed towards oneself and towards the partner may be a correlate of social empathy experienced by individuals who witness their partner under stress. This population-level mirroring of the partner’s emotional response may be critical in motivating a supportive defensive reaction in primates, which buffered both behavioral and neural responses to threats.

## Discussion

The neural mechanisms underlying natural social behaviors in humans and non-human primates have been notoriously difficult to study in ecologically-valid environments. Here we used an ethological approach, new wireless neural data-logging technology, and computer vision techniques to investigate how cortical activity relates to natural behavior in freely-moving, socially-interacting macaques. We manipulated social contexts and exposed monkeys to social threats to investigate the neural basis of affiliative and defensive behaviors, including empathy and social support. We found that activity of units in both prefrontal and inferotemporal cortices was broadly modulated by a wide array of behaviors, with affiliative behaviors characterized by global suppression and aversive behaviors by global excitation. Furthermore, firing rates of individual neurons were modulated by a vast array of behaviors, more than what is typically observed in laboratory tasks, suggesting broadly distributed rather than sparse neural coding of natural behaviors in cortex. The presence and identity of surrounding monkeys strongly structured neural population coding of behaviors, and both behavior and social contexts were highly decodable from population activity. Monkeys spontaneously engaged in remarkably reciprocal grooming interactions, which were unlikely to occur without a form of mental accounting. Neural population activity tracked net number of grooming bouts given and received, as well as elapsed time, a potential mechanism supporting “calculated reciprocity”^53,54^. Threats from a human intruder provoked behavioral agitation and strong neural responses in subjects, both of which were mitigated by the presence of a partner. Neural responses were similar when the subject monkey or his partner were threatened, a neurophysiological ‘mirroring’ consistent with empathy. Importantly, neural signatures of behaviors and social contexts were irreducible to measured visuo-motor contingencies.

Contrary to expectations, populations of neurons in TEO and vlPFC carried comparable information about behavioral states, social contexts and records of past interactions. Given the anatomical and functional distinctions between these areas, it seems likely that the modulations we observed occur in many other cortical areas as well, although it remains possible that areas not recorded here may show more specialized activity during natural behavior. Our findings extend prior work in rodents showing brain-wide behavioral state modulations as a function of movement or arousal in primary sensory areas^45,46,60,61^. Multiplexing of neural representations of behavior and social context in a visual processing like TEO may be a mechanism by which the brain contextualizes sensory inputs to facilitate generation of adaptive behavior. The origins and implications of apparently redundant neural signals across brain areas needs further investigation^62,63^. It is important to note that neuronal activity patterns in TEO and vlPFC may be distinct at shorter timescales than analyzed in the current study. For example, neural responses in TEO may precede those in PFC, but feedback signals from PFC to TEO may blur any differences when data is analyzed at the level of seconds. We advocate application of machine vision-based behavioral quantification, which offers millisecond-scale resolutions, to dissect neural signatures of natural behavior in future studies^64^.

Reciprocity is a fundamental mechanism by which numerous mammals and birds build and maintain strong social bonds^4^. While reciprocity is common in highly social vertebrates, the mechanisms by which it is achieved may vary across species from simple heuristics to cognitively demanding mental accounting of ‘goods and services’ given and received^5^. Our findings revealed that, when interactions were limited to one partner, rhesus macaques kept close track of both the number of grooming bouts and time given and received over days–a cognitively demanding form of mental accounting^6^ which may be afforded by greater encephalization in primates^7^. To overcome some of this computational complexity, detailed mental accounting of past interactions uncovered here may be engaged for a limited number of important partners, whereas simpler heuristics may be used to track interactions within the larger group. Neural mechanisms supporting alternative forms of reciprocity may rely on subcortical brain areas common to all mammals^8^ instead of the cortical representation observed here.

Here, we adopted a neuroethological approach based on the premise that a species’ neural circuitry is refined by evolution to support specific functions with adaptive significance in the natural world^2,67,68^. By letting our subjects behave spontaneously in an enriched social environment, we uncovered neural signatures of an unprecedented array of natural behaviors which would otherwise remain undiscovered through traditional approaches using highly-controlled behavioral tasks^2,69^. We report widespread, highly-structured neural signatures of natural behaviors and social contexts that permeate the day-to-day lives of primates. The highly-distributed coding we observed contrasts with ubiquitous observations of sparse coding^33^, and this discrepancy may reflect the limited and constrained nature of laboratory tasks as the source of neurophysiological data upon which sparse coding models are based. Foundational theories of brain functions may need revision as neurobiological investigations are extended to the complexity of natural behavior. Finally, enabling macaques to express their natural behavioral repertoire allowed the neurophysiological study of fundamental, fitness-enhancing mechanisms–reciprocity and support when facing external threats–at the core of social relationships in primates. This work lays the foundations for an ethological turn in primate neuroscience^69^ to unravel the neurobiological mechanisms that support our own rich and complex social lives.

## Acknowledgments

We thank Robert Seyfarth, Adam Lowet, Richard Lange, Merhdad Jazeyari Labs and, in particular, Juan I. Sanguinetti-Scheck, for conceptual, theoretical and methodological advice. We also thank Xiaowei Jiang for help with pose-tracking, and Jordan Matelsky for artwork for the human-intruder paradigm. Finally, we thank our subject monkeys, Lovelace, Amos, Sallyride and Hooke from which all the data included in this paper is sourced.

## Author contributions

Study conceptualization: CT, ST, MLP; Surgical implantation: ST, KG; Data collection: CT, FP; Behavioral labeling: CT, AA, ST; Video motion tracking: FP; Data analyses: CT, ST, RWD, FP; Figure editing: CT, ST; Funding acquisition: CT, FP, ST, MLP; Writing of manuscript: all authors reviewed and approved this manuscript.

## Data and materials availability

Data is available on Open Science Framework platform and code is available on GitHub upon publication at https://github.com/camilletestard/Datalogger

## SUPPLEMENTARY MATERIAL

### Materials and methods

#### 1. Subjects

Four adult rhesus macaques (*Macaca mulatta*) participated in the experiment, two males, from which we recorded neural activity, and their two female enclosure-mates, which served as partners. No neural data was recorded from females. Due to past aggressive interactions between members of each pair, males and females in this experiment were only allowed to be physically in the same space under the supervision of the experimenter, and all interactions were recorded.

Animals were trained using positive reinforcement via clicker training to enter a primate chair (Hybex Innovations, Canada) where we then lowered a custom-built plexi-glass face shield to stabilize the head for up to 5 minutes while positioning the neural data-logger device and/or for implant maintenance. No other head-restraining methods were used for this experiment. After the data-logger was installed, males were reintroduced into their home enclosure in the colony room to begin the experiment. Subject monkeys underwent no further training, and no food or liquid controls were used for this experiment. The entire study, from starting chair-training until all the data was recorded, lasted 6 months (mid-March to mid-September 2021).

#### 2. Surgical procedures

Surgeries were performed by senior research investigator S.T. and veterinarian K.G. Surgical plans were prepared with the help of brain MR images obtained on a Siemens PRISMA 3T scanner. T1-weighted images (MP-RAGE) with and without gadolinium enhancement (Gadovist®, Bayer, Germany) were obtained to reconstruct the 3D cortical surface and vasculature (Brainsight Vet 2.4, Rogue Research, Canada; see **Fig 2A**). Custom Utah arrays (Blackrock Microsystems, UT) were designed on the basis of local morphology in the region of interest for each monkey. The aim was to cover optimally the regions of interest (ROIs: areas 45 and TEO) while avoiding major blood vessels. For this reason, we targeted the right hemisphere for monkey A and the left hemisphere for monkey H. All surgeries were carried out under isoflurane anesthesia and under strict sterile conditions with assistance from experienced veterinary staff. The neural recording implants (Utah arrays) and surgical procedure are consistent with surgeries FDA-approved for use in human patients. The animal’s head was positioned in an adjustable holding frame (Jerry-Rig, NJ) and a midline skin incision made to expose the dorsal aspect of the skull. The temporalis muscles were retracted and a square-shaped craniotomy was made over the ROIs, based on MRI coordinates, with a dental drill equipped with a diamond round-cutting burr (Horico, Germany). The dura-mater was exposed and a dural flap was performed extending ventrally. Direct visualization of cortical landmarks, including the arcuate sulcus and the sulcus principalis (area 45), and superior temporal sulcus and inferior occipital sulcus (area TEO), was used to determine implantation sites. Arrays were carefully positioned over the ROIs and implanted using a pneumatic inserter held by a flexible surgical arm. The dural flap was closed with 5-0 Vicryl® sutures. The bone flap was thinned with a drill and replaced over the craniotomy and secured to the skull with low-profile titanium plates and screws (DePuy Synthes, IN). A Cereport connector was secured dorso-caudal to the skull opening using eight 1.5mm diameter titanium screws with a length determined by the skull thickness as measured by pre-operative MRI. The two reference wires were inserted between the dura and the cortex and the exposed portion of those wires and of the array wire bundle were coated with a thin layer of Geristore (Denmat, CA). The muscle, fascia and skin were closed in anatomical layers with absorbable 3-0 Vicryl® sutures and the Cereport connector protruded through a small opening in the skin. Animals received cefazolin (225 mg/mL) at 20 mg/kg IV intraoperatively q 2 hrs, and 20 mg/kg ceftiofur CFA (Excede(R), and 100 mg/mL) SC once post-operatively. The anti-emetic maropitant (Cerenia(R), 10 mg/mL) was administered at a dose of 1 mg/kg SC pre-operatively. Analgesia consisted of buprenex 0.02 mg/kg IM once pre-op, then BuprenorphineER (ZooPharm(R), 10 mg/mL) 0.15 mg/kg SC once at extubation. Anti-inflammatories/analgesics: meloxicam 0.2 mg/kg at surgery and 0.1 mg/kg SID x 4 days. Dexamethasone SP 0.5 mg/kg BID on day of surgery. Monkeys recovered for two weeks before the first recording session. No headposts were implanted for this study and no acrylic or dental cement was used (see **Fig S1C** for surgical outcomes).

#### 3. Neuro-behavioral recordings in the home enclosure

##### a. *Setup and behavioral manipulations*

We recorded neural activity from freely-moving, spontaneously behaving rhesus macaques in their home enclosure. This protocol minimized disruption of monkeys’ daily routines, reducing stress as well as training time, thereby promoting expression of natural behavior. The two pairs were located in neighboring enclosures (i.e., side-by-side). We used fully-occluding black room dividers (Versare MP 10 Economical Portable Accordion Partition) to visually isolate our subject monkeys from the other monkeys in the colony room, which contained a total of 15 male and female rhesus macaques. Subject monkeys could hear but not see the other monkeys in the room. Vocalizations and other sounds emitted by non-experimental macaques were captured through microphones but not analyzed for this study.

Each session began with a 15 min installation phase in which we: 1) Installed 8 battery-powered GoPro cameras (HERO7) around the subject’s home enclosure and the neighboring one at locations optimized to monitor all subject monkeys’ behavior (see **Fig 1A** for camera positions); and 2) transferred the subject monkey from his enclosure into a primate chair to install and secure the data-logger. Once the data-logger was secured to the head, the subject monkey was returned to his enclosure and simultaneous neural and video recordings began. Both steps were done in parallel by two experimenters to reduce preparation time. The two human experimenters were seated in front of the experimental enclosures and remained visible to the subject monkeys during the entire length of the session.

Each recording session lasted 2.5 hours and was divided into three experimental blocks, in which we varied social context. There were two 45 min “paired blocks,” in which the male subject was paired with his female partner, and a 1 hour “alone block,” in which the subject was fully visually isolated from all other monkeys. During paired blocks, subjects and their female partners had access to the full enclosure space (height: 2.5 m, width: 2 m, depth: 1 m) and were free to interact. Subject monkeys also had visual access to the adjacent enclosure where only one monkey, dubbed the “neighbor”, was visible at any given time. Subject and neighbor monkeys could not touch each other but could observe each other through the caging. The neighbor monkey was either the male or the female from a different male-female pair. Neighbor identity was switched for each of the two “paired blocks” (e.g., if the neighbor was male in the first “paired block”, the neighbor was female in the second). During “alone blocks”, the subject only had access to half of the enclosure space (height: 2.5 m, width: 1 m, depth: 1 m) and no visual access to other monkeys. **Fig 1A** provides a schematic of the experimental setup and block design. Block order was varied pseudo-randomly across sessions (see **Appendix 4** *“Intervention schedule”* for more details).

Within each block, we performed two additional manipulations. First, we provided naturalistic foraging material at the beginning of each experimental block. We used artificial turf foraging trays (© Otto Environmental) in which we embedded a variety of small food items (on average 300 small items, <50g of rewards per session). Food items included pieces of dried raisins and other berries, small spherical fruit pellets (Bio-serv 190mg fruit pellets), pieces of marshmallow, peanuts, and yogurt drops. By hiding numerous small food items (max 5mm in diameter) inside these foraging trays, we ensured that full depletion of the tray required at least 10 min of continuous foraging. Subject monkeys foraged in multiple bouts throughout sessions (**Fig 1B**). Successful foraging required precise motor coordination to grasp the small hidden items, akin to precision grasps used for grooming.

Second, we performed human intrusions to elicit aggressive and/or defensive behaviors from our subjects, who otherwise would very rarely exhibit aggressive behaviors towards other experimental monkeys. Human intrusions were 30 second-long interventions in which the same male experimenter approached the front of the experimental enclosure and continuously displayed open mouth threats and confrontational stares towards either the subject (threat to subject) or his female partner (threat to partner) (**Fig 5A**). Open mouth threats are common aggressive displays in rhesus macaques, and human threats elicited strong responses from the subject monkeys (irrespective of whether the target was the subject or his partner, **Fig 5B**). Within any given session we performed 6 human threat interventions, 3 towards the partner and 3 towards the subject, across the three experimental blocks. Threats were either performed while the subject and partner monkeys were paired or when the subject was alone to assess the effect of social context. For more details refer to the intervention schedule in **Appendix 4**.

At the end of each recording session, there was a 15 min breakdown phase in which we transferred the subject monkey into the primate chair, stabilized his head using the plexi-glass face shield to remove the data-logger, and performed routine cleaning of the implant by applying alcohol over and around the connector, flushing with distilled water and then with chlorhexidine. Once the connector was fully dried, we placed a protective cap over it and transferred the animal back into his home enclosure. We removed all cameras and occluding panels to return the colony room to its baseline state at the end of each session. We then proceeded to upload the video and neural data collected to a centralized server (Dropbox©).

##### b. *Neural recordings and spike detection*

We used chronically-implanted Utah arrays for neural data recording of single neuron spiking activity. Each monkey was implanted with two 64-electrode arrays, one in frontal cortical area 45 and one in temporal cortical area TEO. Monkey A was implanted in the right hemisphere and monkey H in the left hemisphere to avoid major vasculature on the ROI (**Fig 1A**). The two arrays were coupled to a single Cereport connector with 128 channels. We used a novel custom adapter to connect the Cereport to a neural data-logger to enable range-free, wireless chronic recordings (Spikelog128, Deuteron Tech., Israel). The raw signal was bandpass-filtered (500 Hz to 7000 Hz) and digitized (16 bits) at 32,000 samples per second on the head mounted data-logger. Digitized data was saved on a microSD card (Kingston, 512Gb) inside the data-logger. This allowed for a light (33 g), miniaturized (31×32×27mm), fully wireless neural recording system, enabling the animal to express unconstrained natural behaviors (**Fig S1C**). We used a tablet (Surface Pro 7, Microsoft Inc.) to wirelessly communicate with the data-logger using the Deuteron software via a radio-transceiver box (ST4). During each session, we sent event-markers for behavioral and camera recording events through radio transmission. Event-markers were synchronized to the master clock of the neural recording system.

The Deuteron logger we used amplified, filtered, digitized and stored neural data directly on the head of the animal, within 4cm of the actual recording site. This minimized the effect of environmental electro-magnetic noise, and abolished any noise inherent to wireless transmission of data. Moreover, during surgical implantation of the array, the two reference wires were embedded within the Utah array wire bundle, and protruded immediately adjacent to the implanted array, minimizing the distance between the recording electrodes and their reference. This was crucial to eliminate electromyographic (EMG) noise that interferes with other recording techniques. Moreover, the signal-to-noise ratio of Utah arrays used in this study is known to decrease over a period of months, and along with it the number of individual neurons that can be detected. To benefit from the optimal signal-to-noise ratio window these arrays can offer, our recording sessions occurred within 2 weeks to 2 months post implantation. During that period, the signal to noise ratio we obtained with our recording apparatus was < +/-15uV. Therefore it was possible to observe and isolate single neurons with very low spiking amplitude of less than +/- 20uV (**Fig S1D**).

The entire spike sorting process was done within Plexon Inc.’s latest Offline Sorter (v4.7) software using manual sorting techniques. Although semi-automated and fully automated sorting methods are much faster (e.g. Kilosort), we intentionally took the time to manually sort our spikes for a better understanding of the raw data and easier quality-checks. It is also worth noting that Utah array recordings do not benefit from analytical methods used in Kilosort, which depend on signal redundancy across electrode contacts (Utah arrays have a pitch of 400um). Below, we detail our spike sorting pipeline step-by-step:

1. The raw data was filtered using a 4-pole Butterworth high-pass filter with cut-off frequency of 250Hz, in addition to a hardware bandpass filter of 0.3Hz-7000Hz.
2. The filtered data was then thresholded to detect action potentials (“spikes”). In order to remove as much noise as possible, we used a conservative channel-specific threshold of -3.70 SD from the mean of peak heights histogram (default thresholds are set at -3 SD).
3. Remaining high-amplitude artifacts were removed using the “Invalidate Cross-channel Artifacts” option with parameters “ticks = 10 (315usec)” and “% of channels = 50”. This tool automatically removes shared noise across channels (noise appearing on 50% or more of channels).
4. Using 3D plotting (PC1, PC2, Time), we visually confirmed that the noise level throughout the 2.5h-session for every channel was constant (**Fig S1D**). If it was not stable, the data on that channel was discarded.
5. Putative units were then selected based on the following criteria:

a. The unit had to be present throughout the session. If the unit suddenly appeared or disappeared because of changing isolation, it was discarded.
b. The units’ waveform clusters needed to be segregated based on the first two principal components, as well as non-linear energy. Clusters were manually circled in the 2D-view (**Fig S1D**).
c. The multi-unit cluster of every channel satisfying those criteria was always included as a unit. Hence, every stable channel has at least one unit on it (hence, minimum unit count is 128 per session, assuming all channels are valid).
6. Well-isolated single units as well as multi-unit clusters were manually classified on each channel and saved for future analyses in MATLAB (MathWorks, MA). Example raw data and spike sorted in Offline Sorter can be found in **Fig S1D**.

Based on the assumption that entirely different neurons were recorded in each session, we recorded a maximum of N = 3,689 units (including single and multi-units) over 12 sessions (6 per monkey, mean of 307.42 units per session, SD=19.67) during 1 month of recording. Neuron count remained stable throughout the duration of recordings (**Fig S1E**). Two additional sessions were recorded but the neural data was invalidated due to liquid insertion in the datalogger causing large amounts of noise in the signal. These sessions were kept for behavioral analyses.

We emphasize that we attribute little importance to the number of units recorded in our study and acknowledge that different sorting techniques will yield different neuron counts. An inflated number of recorded units is only a problem if results reported rely on statistical tests where sample size depends on the total number of units across sessions. We did not run any statistical tests in this fashion and no conclusion relies on the total number of units. Therefore we are confident the total number of units does not impact our results or conclusions in any way.

To show that our main results were robust to the spike sorting approach chosen, we re-sorted two sessions (one from each monkey) using more stringent criteria: (1) Multi-unit clusters were excluded entirely; (2) Only single unit clusters without any overlap in PCA space and with an interspike interval larger than 2 ms were included. Using these more stringent sorting criteria, results remain unchanged (**Fig S2E**).

##### c. *Behavioral monitoring*

Prior to the start of the experiment, we conducted 10 hours of preliminary observation of the subject monkeys (5 x 2-hour sessions, 3 sessions for monkey A, 2 sessions for monkey H) to establish a behavioral ethogram. Our group has conducted extensive field research on a free-ranging colony of rhesus macaques on Cayo Santiago island, Puerto Rico, for over a decade (ethogram for Cayo macaques in **Appendix 1**), and we leveraged this experience to devise an ethogram for our study. Our final ethogram included 24 behaviors that recapitulate behaviors observed by our team in the free ranging population, including behaviors such as “foraging”, “groom partner”, “getting groomed”, “lip-smacking”, “submission” and “mating” (see **Appendix 3** *‘Ethogram’* for a complete list of behaviors and accompanying detailed descriptions). The Deuteron custom software allowed us to send event-markers through radio-transmission during recordings. During the experiment, a human observer (the same across all sessions) time-stamped behaviors of the subject monkey, marking the start and end of behaviors from the ethogram, as ethologists routinely do when observing primates in the wild using continuous focal animal sampling^70^.

To complement human-based live observations of natural behaviors, we also video-recorded the behavior of the subject monkey, its partner, and neighbor using 8 GoPro HERO7 cameras (1080p resolution, 29.97 frames per second). Cameras were synchronized to each other and to the master clock of the neural recording system using a GoPro Smart remote via bluetooth with sub-second temporal resolution (<0.5 sec, benchmark established through extensive testing by the authors prior to the start of the experiment). To ensure accurate synchronization, we performed two additional synchronization checks during each session, using an auditory signal captured by the cameras and the Deuteron event log. Cameras were positioned to ensure maximum coverage of the subject’s home enclosure, his partner and the neighbor.

To ensure second-resolution accuracy, behavioral events timestamped live during the session were adjusted post-hoc through manual video scoring by two expert human observers (CT and AA) with extensive primate field observation training. Inter-observer reliability over 3 jointly-scored sessions was 93%. The remaining 11 sessions were coded by only one of the two expert observers (AA) who was blind to all subsequent neural data analyses. We labeled the behavior of the subject, partner, and neighbor to the second resolution. Only the behavior of the subject was related to neural activity for this study; the behavior of the partner monkey will be the focus of future work.

#### 4. Description of behavior

##### a. *‘Rest’ as a reference state*

We categorized timepoints when monkeys did not engage in overt behavior (as defined by our ethogram) as ‘rest’. Rest typically manifested as a monkey sitting and looking around, and thus should not be confused with sleeping. Our subjects were fully awake and behaving for the entire length of the experimental sessions. In subsequent analyses we used rest as a reference state, unless specified otherwise.

##### b. *Comparison with ethogram used with Cayo Santiago free-ranging macaques*

Our laboratory setup allowed our subject monkeys to express a breadth of natural behaviors which largely overlaps (84%) with the breadth of behaviors observed in a free-ranging colony of rhesus macaques (**Appendix 1**, ^71^). To quantify this overlap, we made a list of all behaviors in the Cayo Santiago ethogram and counted the number of behaviors which were also exhibited in the laboratory (see **Appendix 2**).

*Cayo-Lab Overlap = [# behaviors exhibited in the lab / All behaviors exhibited on Cayo]*

##### c. *Co-occurrence of behaviors*

The subject occasionally engaged in multiple behaviors simultaneously (e.g., “scratch” and “getting groomed”). In these cases, we utilized the following criteria for choosing the behavior to code: When a behavior co-occurred with “proximity”, “rowdy room” (loud noise in the colony room), or “other monkeys vocalize”, we prioritized the former behavior (most common case). In all other cases of co-occurrence of behaviors (< 2% of the time), we established a “hierarchy” of behaviors that take precedence (e.g., getting groomed takes precedence over scratch). See **Appendix 3** *‘Precedence’* for more details. The same hierarchy of behavior was applied to the partner’s behavior.

##### d. *Exclusion of behaviors in analyses*

During behavioral labeling, “proximity”, defined as being within arm’s length of another monkey, was coded as part of the ethogram. In subsequent analyses, we pooled “rest” and “rest in proximity” because we judged the recording arena not large enough to make a meaningful distinction between them (paired monkeys were essentially always in proximity). The majority of our analyses used “rest” epochs as a reference and excluded behaviors performed by other monkeys in the colony (e.g., “Other monkeys vocalize” and “Rowdy Room”, see **Appendix 3** for definitions), and rare behaviors (which happened for a duration of <30 s in a given session, or < 10 times across all sessions, see **Appendix 3** *‘Behavior lists’* for a list), unless specified otherwise. These exclusion criteria resulted in the majority of our analyses considering 17 of the 24 total behaviors observed across sessions (see **Appendix 3** *‘Focus of analyses’*). Typically, 12 out of these 17 behaviors occurred frequently enough to be analyzed in any given session.

##### e. *Behavioral frequency and transition matrix*

We combined behavioral data from all sessions and computed the proportion of time spent in each behavioral state: proportion for behavior A = [# of seconds in behavioral state A / total # of seconds] (**Fig 1C**). We then computed behavioral transition matrices for each monkey in which each cell is the total count of transitions from the row behavior (y-axis) to the column behavior (x-axis) across all sessions (**Fig S1B**). Transitions excluded “rest” epochs, behaviors performed by other monkeys in the colony (“Other monkeys vocalize” and “Rowdy Room”), as well as short event behaviors (1-3 s in length: “vocalization”, “lip smack”, “anogenital investigation” and “scratch”). Transition plots are pooled across monkeys and thresholded by frequency per behavior in **Fig 1D** (>20% transitions), and un-thresholded in **Fig S1B**.

##### f. *Grooming categories*

It is possible that neural responses to a behavior could be modulated by preceding events. In our experimental sessions, we were able to identify three grooming contexts that are known to be important from decades of field observations of primates in the wild ^72^: 1) Whether grooming was initiated by the partner or the subject; 2) Whether a grooming bout immediately followed a previous one, i.e. reciprocal grooming and 3) Whether grooming occurred after aggression (see **Appendix 3** *‘Grooming categories’* for detailed description). Note that these categories are not mutually exclusive. For example, a grooming bout can occur after aggression and also be initiated by the subject.

##### g. *Grooming reciprocity*

Grooming reciprocity was calculated using two metrics: reciprocity (1) and directional reciprocity (2).

(1) Reciprocity = 1 - abs(Groom given - groom received)/Total grooming. Values will range between 0 –fully one-sided– and 1–perfectly equal.
(2) Directional reciprocity = 1 - (Groom given - groom received)/Total grooming. Value will range between 0 and 2, where 1 = perfectly equal, <1 more was given than received and >1 more was received than given.

Note that grooming duration given vs. received or number of grooming bouts in each direction could be used to compute reciprocity. We refer to the former as “groom duration reciprocity” and the latter as “bout number reciprocity”.

To investigate reciprocity in grooming bout number and duration between members of a pair, we considered unidirectional grooming bouts separated by less than 10 seconds to be part of the same bout. Alternating bouts consisted of grooming bouts preceded by grooming in the opposite direction (Give - Receive - Give). When examining the distribution of inter-bout intervals (IBI), we identified a bi-modal distribution with very short IBI and much longer ones (**Fig S4C**). We defined alternating bouts separated by <20 sec as “turn-taking”.

To characterize how reciprocity developed and fluctuated over the course of a session, we computed a running reciprocity metric that considers all grooming seconds and bouts up to the current time in the session (**Fig 4D**). In all sessions grooming occurred for at least 2500 sec (some sessions extended grooming for longer). Therefore, to compute average running reciprocity across sessions, reciprocity was computed for the first 2500 sec of grooming within a session and aligned to “Start of grooming”.

#### 5. Simulations for grooming reciprocity

Simulations were used to establish the distribution of reciprocity and turn-taking assuming monkeys groomed at random (null distribution). If the observed reciprocity and turn-taking fell outside this distribution (*P* < 0.05), we concluded it was unlikely monkeys groomed randomly and instead surmised grooming decisions were made to reciprocate and/or take turns. All simulations were run 10,000 times to obtain p-values.

##### a. *Reciprocity in bout number given random allocation of grooming*

To generate a null distribution for bout number, we chose to model grooming as the combination of an initiation process and a duration process. At each moment a monkey can choose to initiate grooming his partner or not. We modeled this choice by using a random draw from a uniform distribution to set the probability of choosing to start grooming and then taking a random draw from a Bernoulli distribution with that probability (i.e. “flipping a coin” with probability *p* of initiating grooming). If the monkey did not initiate grooming, we conducted the same process to see if his partner would initiate grooming. If neither monkey initiated grooming, we moved on to the next moment and repeated the simulation. If either monkey initiated grooming, we took a random draw from an exponential distribution with a expected duration of 100 seconds and set the duration of the initiated bout to this random draw. We continued the simulation until 45,000 seconds (approximately the length of all sessions combined for one monkey) had elapsed from time waiting for either monkey to initiate grooming and total duration of the grooming bouts. We then tallied the number of grooming bouts for each monkey and calculated the reported statistics in **Fig S4A** top panel. Note that the range of total number of grooming bouts from this simulation matched the observed total number (range: 153-231 bouts).

##### b. *Probability of turn-taking given random ordering of bouts given and received*

To obtain a null distribution of turn-taking, we shuffled the directionality of observed grooming bouts (i.e. give vs receive). Bouts that alternated in direction and were separated by less than 20 sec were labeled as “turn-taking” in the shuffled data. We then calculated the proportion of turn-taking bouts out of all grooming bouts and compared this null distribution to the observed proportion of turn-taking (**Fig S4A** middle). Finally, we calculated the probability of observing the true proportion of turn-taking in this null distribution (p-value).

##### c. *Reciprocity in groom duration given random allocation of grooming length to observed grooming bouts*

We generated a null distribution of reciprocity in groom duration assuming monkeys did not track time. We maintained the empirical frequency, timing and directionality of bouts but randomly sampled durations from the observed distribution of grooming bout durations (**Fig S4D**). This way we kept the natural statistics of the grooming behavior of our subjects. We then compared this null distribution to the observed reciprocity values, both within and across sessions, and calculated the probability of observing the observed reciprocity (**Fig S4A** bottom).

#### 6. Behavioral quantification

##### a. *Monkey detection and pose estimation*

To detect monkey’s position and estimate their pose, we trained a monkey detection model and a pose estimation model using the *mmdetection* and the *mmpose* Python libraries (Python version 3.8) ^73,74^ respectively. Combining these models ultimately enabled us to quantify the position of 17 body landmarks in freely-interacting monkeys (see **Appendix 3** *‘Visuo-motor measures’* for full list) by following the training procedure we will now outline. For both models, we leveraged standard data augmentation techniques as well as transfer learning ^75^. We first trained each model on the MacaquePose dataset ^76^, and then supplemented training with annotated images from our dataset. We trained the monkey detection Faster R-CNN model ^77^, and then the pose estimation model HRNet-W48 ^78^, which has been shown to perform well in tracking myriad primate species ^79^. Pretrained models for both the Faster R-CNN and the HRNet-W48 are publicly available for training in the *mmdetection* and *mmpose* Python libraries respectively. For the monkey detection model, we labeled and split 3,082 images (3,962 monkey instances) of freely-interacting monkeys into training (70%), validation (15%), and testing (15%) sets using Roboflow ^80^. For the pose estimation model, we labeled and split 4,889 images (6,084 monkey instances) into training, testing, and validation sets using the Scikit-learn Python package ^81^. We used mean average precision (mAP), precision, and recall metrics to evaluate the monkey detection model, and mean average precision to evaluate the pose estimation model. The trained detection model returned a mAP of 99.4%, precision of 98.5%, and recall of 99.3%, and the trained pose model returned an mAP of 93.2% across body landmarks. After training these two models, we analyzed the video data using a three-stage pipeline: 1) detect monkeys, 2) assign their bounding boxes IDs and 3) quantify their poses. Detection of the monkeys was handled directly by the monkey detection model. The subject monkey (ID 1) and partner monkey (ID 2) bounding boxes were assigned based on camera angles during the "alone" block, which we verified manually. If the camera angle faced the subject monkey during the alone block, we assigned all bounding boxes during this block ID 1, and if it faced the partner monkey, we assigned all bounding boxes during this block ID 2. During social blocks, we employed a set of heuristics that relied on the presence of a data-logger to assign IDs to the bounding boxes of detected monkeys. Namely, we assigned IDs to monkey bounding boxes based on the top center point of each bounding box and its proximity to the data-logger bounding box top center. For frames in which both monkey bounding boxes and data-logger bounding boxes were detected, we computed the top center point of each bounding box and assigned the monkey bounding box "1" to the monkey bounding box top center that was closest to the data-logger bounding box top center. For frames in which the data-logger was not detected, we used previous bounding box data to assign IDs based on closest distance to bounding boxes in previous frames. Once the IDs were assigned, we performed pose estimation on the monkeys within each bounding box. To smooth the pose tracking results, we applied a Savitzky-Golay filter consistent with prior primate pose estimation studies ^82^. Of the 7 cameras surrounding the monkey enclosure, we analyzed behavior from the side angles (four out of seven camera angles), which had the better overall view of the animals’ pose. For each predicted keypoint, the pose estimation model returned a corresponding confidence score as a measure of how likely it is that the keypoint exists in that position. The confidence score is based on the output of a “softmax” activation function that assigns a probability from 0 to 1 to each possible keypoint location in the image; the higher the probability, the higher the confidence score. After inspection, we considered a confidence cutoff value of 0.2 to be appropriate (see sample tracking results in **Movie S5-8**) and conditioned the X,Y coordinates using this threshold for future analysis (i.e. all tracking data with a lower confidence score than our cutoff value were discarded from downstream analyses).

##### b. *Field of view quantification*

To quantify the field of view of each subject monkey, we used the pose tracking software DeepLabCut to track the top and bottom positions of the data-logger (DeepLabCut version 2.2.2; ^83,84^). We labeled 6,486 images (2,423 monkey instances) taken from top-down view videos across all 12 sessions and used 95% of these images for training. We trained a ResNet-50 neural network to perform pose estimation with default parameters for 2 million iterations, achieving a mean average Euclidean error (MAE) training error of 2.66 pixels and MAE test error of 5.54 pixels. Like the previous pose tracking model, this neural network outputs a likelihood score for each predicted keypoint. As a confidence metric, we used a p-cutoff, a parameter that determines the confidence threshold of each keypoint, of 0.7 to condition the X,Y coordinates for subsequent analyses. After training, we used this model to track data-logger position from the held out sessions. Using the front and back corner coordinates of the data-logger, we calculated the angle, in degrees, between these two points using the *atan2d* function in Matlab. The back corner of the data-logger was used as a reference point around which the front corner rotates. This rotation angle is equal to the orientation of the head in space, which we termed “field of view”. Sample tracked results are shown in **Movie S9**.

##### c. *Metrics and missing data for movement and field of view tracking*

For each session with appropriate side and top-down view videos (used for full body and logger tracking, respectively), we extracted various visual and motor metrics, including animal’s x,y position, full body key points, head movement, velocity, acceleration, and head direction (see **Appendix 3** *‘Visuo-motor measures’* for a full description). We had to exclude 2/12 sessions for analyses including full body tracking due to missing side view videos on these sessions.

Our subject monkeys sometimes were out-of-sight of camera coverage leading to some key points becoming invisible for periods of time. When these moments occurred, our tracking models attempted to infer the position of invisible body-parts based on a full skeleton model of the animal. It was not always possible for our models to make a high certainty inference; therefore there is data missing from our movement tracking. Using a 0.2 confidence score for pose tracking with *mmdetection* and *mmpose* and 0.7 score for data-logger tracking with DeepLabCut, we had 78.98% missing data on average across sessions (SD 28.34%). We omitted these time-points from subsequent analyses.

##### d. *Manual scoring of agitation during human threat paradigms*

We manually scored the behavioral response to human threats from video recordings by considering the occurrence of three types of social behaviors by the subject monkey: 1) attacks: defined as rapid arm gestures by the monkey aimed towards the human intruder, 2) dodges: defined as rapid head and body movements to achieve an aggressive posture 3) submissions: defined as a presenting hindquarters to the human intruder. The summed frequency of these three behaviors during a single 30 sec human intruder bout was considered the “agitation score” for this bout, such that a high score corresponds to a high agitation level and a low score low agitation. Each bout was classified as either belonging to the “alone” or “paired” category, depending on whether the monkey was alone or paired during the bout. The same scoring procedure was repeated during threats directed toward the partner monkey (**Fig 5B**).

#### 7. Preprocessing steps for subsequent analyses of neural activity

For each session, we aggregated the spike counts of each neuron into time bins of 1 seconds except for analyses involving movements where we binned at the camera frame-rate or 29.97 Hz. This process produced N sets of 1 x T spike count vectors where N is the number of neurons and T is the total number of time bins given the temporal resolution. We then convolved these spike count vectors with a 1 time bin standard deviation gaussian kernel to obtain the smoothed approximate instantaneous firing rate of each neuron. All downstream analyses used instantaneous firing rates as input unless specified otherwise. Units that had very low (<0.1Hz) or very high mean firing rate (>150Hz) were excluded from our analyses.

We also created a 1 X T current behavior vector by assigning each time bin a numeric label which corresponded to a particular behavior from our ethogram (**Appendix 3**). Accordingly, these preprocessing steps created a “neural firing matrix” of size [#units X Time (at desired temporal resolution)] and a “label vector” of size [1 X Time (at the same desired temporal resolution)]. All subsequent analyses were performed on this neural firing matrix and label vector. All analyses were performed using Matlab 2021b unless indicated otherwise.

#### 8. Single unit responses across macaques’ natural behavioral repertoire

##### a. *Responsiveness to behavioral events*

We quantified units’ responsiveness to behavioral events by computing a behavior transition index comparing activity 10 sec before (‘pre’) and 10 sec after (‘post’) the onset of a behavioral state (**Fig 2C,D**).

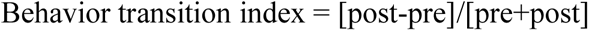

Behavior transition indices range from -1, corresponding to strong suppression following an event onset, to +1, corresponding to strong activation. We focused on state behaviors which lasted more than 10sec and occurred at least 5 times in the session. This narrowed the focus of our analysis to 6 behaviors across sessions: aggression, groom partner, foraging, self-groom, getting groomed and drinking.

We then used two-sample *t*-tests (N *t*-tests= number of neurons x number of behaviors investigated) to determine whether the differences in firing rate between the pre- and post-event epochs were statistically significant. To control for false positives, we used Benjamini & Hochberg’s (1995) false discovery rate (FDR) procedure^85^ as implemented in the Matlab function *fdr_bh*^86^. We considered a unit’s activity to be significantly modulated by a behavior if FDR-corrected *P*-value<0.01. In **Fig 2D** we show behavior transition indices for units with statistically significant modulation comparing pre- and post-event epochs.

Note that representing neuronal responses aligned to specific events in per-event time histograms (PETHs), as in **Fig 2C**, may not be ideal when analyzing neural signatures of natural behavior because precise repetitions of specific event sequences are rare. For example, in session A290721 which provides the basis for **Fig 2C**, the most frequent transition was groom receive to groom give, with 8 transitions (**Fig S1A)**.

##### b. Modulation of activity relative to reference state of ‘rest’

To further explore the response profiles of single units across macaques’ natural behavioral repertoire we extracted the empirical distribution of firing rates of all units during each individual behavior. Each sample in these distributions consists of the firing rate recorded for 1 sec during the session; e.g., if grooming occurred for 1000 seconds during a session, the distribution of firing rate during grooming will contain 1000 data points. We then performed the same operation during “rest” epochs, our reference, and compared this baseline distribution to the distributions we observed for each behavior **(Fig 2E**). Specifically, we used Cohen’s d to quantify the standardized mean difference in firing rate during each behavior compared to rest (**Fig 2F**) via the following formula:

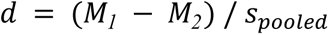

We then used two-sample t-tests (N *t*-tests= number of neurons x number of behaviors) to determine whether the differences in mean firing rate as calculated by Cohen’s d were statistically significant (**Fig S2B**). To control for false positives, we used the same approach as described in the previous section, the Benjamini & Hochberg’s (1995) false detection rate (FDR) procedure. We considered a unit’s activity to be significantly modulated by a behavior if FDR-corrected *P*-value<0.01 (**Fig 2H**). Sample size influences *P*-values, so we repeated our analysis to find the number of behaviors a given neuron is responsive to by subsampling our data to have 100 observations (seconds) per behavior (**Fig S2C**). We also repeated these analyses for well-isolated units only and found qualitatively similar results (**Fig S2D**). To quantify global trends across all neurons we also computed the mean z-scored firing rate of individual units for each behavior and plotted the resulting distribution across all units in **Fig 2H**.

##### c. *Modulation of activity during threats*

To compare the responses of individual units to threats directed towards the subject when alone vs. paired, we computed the mean firing rate of each recorded unit during each condition (threat to self when paired and threat to subject when alone) and plotted one against the other in **Fig 5C**, right panel. Note here that we averaged firing rates across bouts and duration of threat bouts to obtain one value per neuron per condition. Most (71.4%) units showed higher average firing rates to threats when the subject was alone compared with when his partner was present.

To compare neural activity in response to threats towards the subject vs. his partner while they were paired with each other, we first calculated the average neural responses across all neurons in each condition. Here we retained information on variation in activity during a bout and obtained a matrix size [N_neurons x Time] containing the average neural population response across bouts for each condition. For each session, we computed the correlation coefficient between the two average neural population responses during threat to subject compared to threat to his partner while paired. This correlation coefficient is called a “representational similarity score”, as is typically computed in neuroimaging analyses.

#### 9. Linear decoding of behavior and context based on neural population activity

To assess what information a hypothetical downstream brain region could linearly readout from the recorded neural activity, we ran a series of decoding analyses. In each analysis, we used a linear support vector machine (libSVM ^42^) with 5-fold cross-validation. Note that 5-folds were created by splitting the session length into 5 chunks rather than random sampling of indices (which is the default in Matlab but can lead to overfitting). We trained the SVM on simultaneously recorded units, i.e., “the neural population”, to get an estimate of overall linearly decodable information contained in the entire population of recorded units (rather than assessing individual units separately). We used a shuffled, hold-out cross-validation procedure where, for each fold, time points from the session were randomly sampled and assigned to either the training or testing set (training to test split was 80% to 20%). The accuracy of the trained model was calculated based on the number of correct predictions on the testing set (correct predictions / all predictions). There were large discrepancies in the number of observations for each variable we were attempting to decode. To account for how this difference could bias performance, we balanced the number of observations per variable (e.g. current behavior or social context) by randomly subsampling the variables with more observations so that each had the same number of observations as the least observed variable. To account for variance in performance caused by random subsampling, we ran 100 iterations of each SVM and computed the mean decoding accuracy across all iterations. We calculated empirical chance performance of the decoder by randomly permuting the variable labels (shuffled control) before training and following the same analytical procedure.

We ran separate decoders on the neural population to readout information about one of four different variable groups: 1. The current behavior of the animal; 2. Paired vs alone contexts; 3. Neighbor identity when the animals were paired (1-3 in **Fig 3E-F**) and 4. Reciprocity and time elapsed in the session measured during grooming; 5. Time elapsed in the session during rest (4-5 in **Fig S4F**). We used a temporal resolution of 1 second for each decoder.

#### 10. Unsupervised dimensionality reduction of neural population responses (UMAP)

We can treat the activity of a population of simultaneously recorded neurons as a multi-dimensional space where each dimension corresponds to the activity of an individual neuron. A single point in this space therefore would correspond to a particular pattern of neural activity across the entire population and is typically called the neural population state, or neural state for short. There can be meaningful structure in the collection of neural states observed throughout a recording session, and we can look for such structure by visualizing the data with dimensionality reduction methods ^87^. We chose to use the state-of-the-art, non-linear dimensionality reduction technique UMAP (Uniform Manifold Approximation and Projection, ^40^) to see whether meaningful structure was present within our data. UMAP attempts to retain both the global and local structure of the data, such that the overall “shape” of the data is preserved (global structure) and points that were close in the original space tend to remain close in the dimensionality-reduced space (local structure). We used the firing rates in Hz of all simultaneously recorded units during each behavior as observations (excluding rest and rare behaviors) and reduced the data down to three dimensions. We used the matlab implementation of UMAP toolbox ^88^, with parameters: ’n_neighbors’=15, ’min_dist’=0.1, ’n_components’=3. We note that UMAP can be used as a supervised dimensionality reduction technique, but we chose to use the unsupervised variant. This choice ensured that the structure UMAP found was solely due to patterns in the neural states irrespective of information about the current behavior. We plotted the neural states on the first 3 UMAP dimensions and ***post-hoc*** color-coded data points according to what the subject monkey’s current behavior was at that time point (**Fig 3A,B**) or the social context (**Fig 3C,D**). We tried multiple parameter configurations and obtained qualitatively similar results (**Fig S3 E-H**).

We also visualized population activity only considering grooming seconds (**Fig 4K** top panels) or rest seconds (**Fig 4K** bottom panels). Post-hoc, we color-coded each data point by the groom or rest bout number that second belongs to.

#### 11. Calculating distance between neural responses in UMAP space

We can quantify how differently a neural population represents one particular behavioral state or context versus another by comparing the patterns of population activity during each behavioral state/context. One way to perform this analysis is to calculate the pairwise Euclidean distance between the observed neural states (see previous section for detailed definition) for each pair of behavioral states/contexts. Unfortunately, using Euclidean distance can yield surprisingly uninterpretable results in a high dimensional space. In brief, it has been previously shown that using Euclidean distance to calculate the nearest neighbor to a query point in a high dimensional space (i.e., >30 dimensions ^89,90^ can yield the paradoxical result that all data points are equally close or far to the query point (i.e. many/all points are nearest neighbors). We recorded on average 300 units per session and were concerned that we would encounter this dimensionality issue if we used Euclidean distance in the original firing rate space. To mitigate this problem, we first reduced the dimensionality of our data with UMAP (see previous section for more details), an algorithm designed to preserve both local and global structure in the data as much as possible, and then calculated the pairwise Euclidean distance between neural states in this reduced space. We measured distances between neural states (a) across behaviors controlling for the effect of social context (**Fig S3C**); (b) across social contexts controlling for the effect of behavior (**Fig S3D**); (c) during a threat event compared to a reference state of ‘rest’ (**Fig 5C**).

##### a. *Distance between behavioral states, controlling for social context*

To quantify clustering in UMAP space across all sessions, we measured the average distance within and between behavior across all sessions. To do so, we first reduced the dimensionality of the full session’s neural data to 20 dimensions using UMAP (’n_neighbors’, 15, ’min_dist’, 0.1, ’n_components’=20). Then, we measured the Euclidean distance between neural states of different behaviors using the matlab function pdist. For a given session, we subsampled down to 30 time points per behavior and calculated the distance matrix for these points. We repeated this subsampling procedure 500 times and averaged the distance matrix across all subsamples. To compare activity across sessions, we computed the mean distance between neural states within and across behavioral categories during one social context, the female neighbor block (**Fig S3C** is averaged across all sessions). To measure how similar neural population activity patterns were to “threat to subject” (**Fig 5D**), we computed similarity score 1/1+d(pi,pj), where d(pi,pj) is the mean Euclidean distance between two states pi and pj. These results were qualitatively the same irrespective of the number of UMAP dimensions used (n_components=3, 20 or 50).

##### b. *Distance between social contexts, controlling for behaviors*

We used a similar approach than described above to measure distances between states from different social contexts, this time subsampling 100 time points during ‘rest’ across all three contexts (i.e., alone, paired with female neighbor, paired with male neighbor).

##### c. *Distance between a threatened and a baseline state when paired vs. alone*

We considered a baseline state to be the center of mass (i.e., the mean) of all ‘rest’ neural population states. For each human threats towards the subject, which lasted 30 seconds, we extracted a 90sec window: from 30 seconds before threat onset to 60 seconds after. For each second in that window, we computed the Euclidean distance to baseline in UMAP space. We then split threats towards the subject which occurred when the subject was alone to those where he was paired with his female partner. We normalized distances to 20sec prior to threat onset and plotted this normalized distance to baseline in neural firing space in both conditions (**Fig 5C**, lines are mean and shaded areas are SEM).

#### 12. Addressing movement and field of view potential confounds

Allowing animals to move freely enables the expression of natural behaviors but reduces control over visual inputs and motor outputs. Accordingly, we tested whether our results were merely due to the visuomotor experience of the animal rather than their current behavioral state and social contexts. To do so, we ran a series of complementary analyses (note that all analyses in this section excluded timepoints with missing video tracking data). First, we quantified variation in movement patterns and field of view (FOV) within behavior and overlap across behaviors. Second, we measured explained variance in neural activity by movements and FOV compared to behavioral state. Third, we regressed-out movements and FOV and assessed the effect on decoding accuracy of behavioral state. We discuss each analysis in detail in the following subsections:

##### a. *Preprocessing: aligning video and neural data*

We extracted all movement and FOV-related variables from the video data (**Fig 3G**, see video tracking section for more details). For all analyses, we resampled the neural data at 29.97Hz, to match the videos’ temporal resolution. We then aligned neural activity with the onset of video recordings. The neural recording system was always started first and stopped last, such that we timestamped when the cameras turned ON and OFF using the same clock as the neural data. We simply trimmed neural activity which occurred before the cameras were started and after they were turned off. These two pre-processing steps allowed the alignment of movement, FOV, behavioral and neural data required for subsequent analyses.

##### b. *Approach 1: Quantifying variability in animal position, movement and field of view within and across behaviors*

Across all N=12 sessions, we tested whether there was variation in the visual input of the animal across different instances of a given behavior, and whether similar visual inputs could occur during different behaviors. This was done to demonstrate a potential dissociation/decorrelation between behavior and visual input. As a proxy for field of view (FOV), we calculated the head direction of the animal based on automatic tracking of the neuro-logger position in videos (see above for method).

We plotted the distribution of FOV during four different example behaviors and found that each behavior was characterized by a wide distribution of head orientations, encompassing distinct fields of view and, thus, visual inputs during the execution of a given behavior (**Fig 3H**). Invariant behavioral decoding accuracy amidst changing visual inputs rules out the possibility that the neural signature of a given behavior is entirely explained by a specific visual input and suggests the underlying cortical neural representations of behavior are robust to varying sensory environments.

Second, we tested whether the FOV was similar or different across behaviors and during the 3 tested social contexts. If FOV were different across behaviors or contexts, it could indicate a potential confound between the neural signatures of behaviors and visual input. We find similar distributions of fields of view occurred during multiple distinct behavioral states (>50% overlap), further dissociating visual inputs from neural coding of behavioral states (**Fig 3H**). Similarly, fields of view largely overlapped (75% overlap) across different social contexts (**Fig S3K**), yet social contexts were still decoded accurately from neural population activity (see **Fig 3F**). These findings rule out the possibility that a correlation between visual input and context explains all of the neural signatures of behavior and social context observed in both monkeys.

In addition to overlap in field of view, we find similar kinematics across two neurally distinct behaviors, self-grooming and allo-grooming. After quantifying macaque movement with the HRNet pose tracking model (see previous section on video tracking), we isolated the shoulder, elbow, and wrist 2D trajectories of the subject. We then computed the change in shoulder-elbow-wrist joint angle during two 2-min bouts of self- and allo-grooming (**Fig S3J**). It was nonetheless possible to linearly separate the neural representations of these two behaviors (**Fig 3E**), which would not be possible if movement kinematics solely defined these neural signatures.

Taken together, these results show a decorrelation between visual input and behavioral state or context, which weakens the hypothesis that neural signatures of behaviors or contexts in TEO and vlPFC are entirely explained by unique visual inputs or movement patterns.

##### c. *Approach 2: Measuring relative variance explained by movement and field of view compared to behavior*

Next, we ran a multilinear ridge regression analysis to determine how much of the neural variance of each recorded unit could be uniquely explained by the current behavioral state of the animal vs. the sensorimotor environment (**Fig S3L**). The smoothed firing rate for each unit throughout the session was used as the response variable, and the behavioral labels and visuo-motor measurements were used as predictors (see **Appendix 3** for a full description). This method closely replicates the one used in Musall et al. 2019 ^45^ and Tremblay et al. 2022 ^32^ to dissociate the neural variance explained by task variables and movement variables. We used 10 sessions for this analysis, 2/12 had to be excluded due to missing side view videos used for pose-tracking (see above for more details).

Predictors could be associated with multiple regressors in the analysis and were labeled as either analog predictors or event predictors (**Fig S3L**). Event predictors had distinct onsets and offsets and could be represented in the model by a regressor that takes on value of 1 when the predictor was occurring and 0 when the predictor was absent (e.g., the foraging behavioral state). To account for preemptive or lagged neural responses, we constructed “event kernels” for each event predictor. Event kernels were comprised of a set of regressors each of which represented a particular delay from the event onset. We included delays from one second before to one second after event onset. It can be useful to think of event kernels as related to the more familiar peri-stimulus time histogram (PSTH). That is, imagine each delay regressor corresponds to one bin of the PSTH; thus, the entire set of delay regressors allows the model to fit a PSTH to each event predictor. Analog predictors, by contrast, took on continuous values throughout the session and could be represented in the model by a regressor that tracks the measured value (e.g., current head position). We noted that it was possible that neuronal responses corresponded with certain analog predictors crossing particular thresholds rather than corresponding with the exact value of that predictor, and, furthermore, that the responses to these threshold crossing could be preemptive or lagged. For example, a neuron could increase its firing in some stereotyped way before an animal initiates a large arm movement. To account for such responses, we created event kernels for the relevant analog predictors. We used a threshold of two standard deviations to determine what qualified as a significant threshold crossing event for an analog predictor and constructed the event kernel as above. We collected all of the regressors resulting from the above processes into an *Observation* X *Regressors* matrix while keeping track of which sets of regressors were associated with which predictors. This matrix was used to fit a linear model with ridge regression regularization in all subsequent analyses.

To determine the proportion of variance explained by each predictor, we used a 10-fold shuffled, hold-out cross-validation procedure to obtain the predicted neural activity by the linear model for each recorded unit individually. We then calculated the correlation between the predicted neural activity and the actual recorded neural activity for all neurons recorded during a session, yielding a single, overall *R*^2^ for the neural population. This *R*^2^ value gave us an estimate of the variance explained by our model when using all possible predictors. To compute the variance explained by a particular predictor group (e.g. Movements), we refit the model but shuffled all other predictors except the predictor of interest, which we called the “Full contribution” model. The explained variance by each “Full contribution model shows the maximum potential predictive power of the predictor of interest (e.g. Behavior ID). To measure each predictor’s *unique* contribution to the model, we instead only shuffled the predictor of interest and kept all other predictors as they were, which we called the “unique contribution” model. The difference in explained variance between the model using all predictors and this model yielded the *Unique contribution* (Δ*R*^2^) of that predictor. The unique contribution shows the amount of explained variance in the full model that can be uniquely attributed to the group of model variables associated with that predictor (i.e. nonredundant to all other predictors). To account the potential of overfitting we included a ridge regularization penalty in the regression model. Briefly, ridge regularization effectively sets a limit on the sum of the squared regression coefficients which in turn drives regressors that are not contributing much to model predictions toward, but not to, zero. The strength of this regularization is determined by a scaling parameter called lambda. In our analyses lambda was set by cross-validation before the model was fit and the same lambda was used for the full and unique contribution model. Again, for more details, see Tremblay, Testard, *et al.* 2022 as well as Musall *et al.* 2019 ^45^.

##### d. *Approach 3: Regressing out variance explained by movement and field of view for decoding*

To further investigate whether neural activity patterns were merely explained by the immediate sensorimotor environment, we re-ran our decoding analyses of the behavioral ethogram after removing neural variance explained by movement and field of view (**Fig 3J**). To do so, we ran a multilinear regression with neural activity as a response variable and sensorimotor variables as predictors, and used the raw residuals of this regression as input to downstream analyses (see Methods section on decoding above for more details). Residuals of this regression correspond to movement and FOV-controlled residual neural activity and the decoding was run on these residuals. We also ran a version of the decoder where residuals from a regression run between neural activity and behavior ID were used. This decoder effectively attempts to decode behavioral states from visual and movement inputs alone. Since behaviors are largely defined by stereotypical movement patterns, decoding accuracy for behavioral state did not fall to chance, as expected. These two decoding analyses allowed us to measure the information content of the neural population activity about behavioral state, irrespective of visuomotor confounds.

#### 13. Predicting reciprocity from neural population activity using ridge regression

To investigate whether neural population activity in TEO and vlPFC tracked how reciprocal grooming interactions were at any given second in a session, we used multi-linear ridge regression analysis. We tested how the neural population activity predicted two reciprocity metrics and elapsed time (i.e. response variables, see cartoon in **Fig 4G**): (1) net groom duration (Groom duration received - duration given) and (2) net number of groom bouts (Groom bouts received - bouts given) (3) Time, a continuously monotonically increasing variable. Predictors were the neural population activity (i.e. Time (s) x Neuron matrix). Both predictor and response variables were z-scored. We focused our analysis, first, on grooming time points exclusively, where we were most likely to predict grooming reciprocity from the neural activity. Grooming bouts which occurred for 5 sec or less were excluded from this analysis.

To determine model fit, we used a 5-fold hold-out cross-validation procedure to obtain the predicted reciprocity metric by the linear model from the whole neural population. Note that each fold corresponded to a continuous chunk of data in the session, rather than random sampling of indices within the session, which can lead to overfitting. We then calculated the correlation between the predicted reciprocity metric and the actual metric during a session, yielding an overall *R*^2^. This *R*^2^ value gave us an estimate of the variance explained by our model when using all simultaneously recorded neurons (**Fig 4H**). To investigate whether any given single neuron activity tracked reciprocity, we repeated the aforementioned analysis but for each neuron individually (**Fig S4I**). Finally, we asked how well elapsed time could be predicted from the neural activity outside of grooming. We repeated the ridge regression analysis described above but now only including data from non-grooming time points (**Fig 4J**).

## SUPPLEMENTARY FIGURES AND TABLES

### List of Appendices

1. Cayo ethogram, basis for the experiment’s ethogram
2. Comparison of Cayo and Lab ethogram
3. Ethogram (description of behaviors, grooming contexts, visuomotor variables)
4. Intervention description and schedule

### Supplementary Movies

1. Movies S1-4: Example clips of natural behavior in the lab
2. Movies S4-8: Full body tracking certainty cut-off 0.2
3. Movie S9: Logger tracking at 0.7 cutoff.

**Fig S1.**
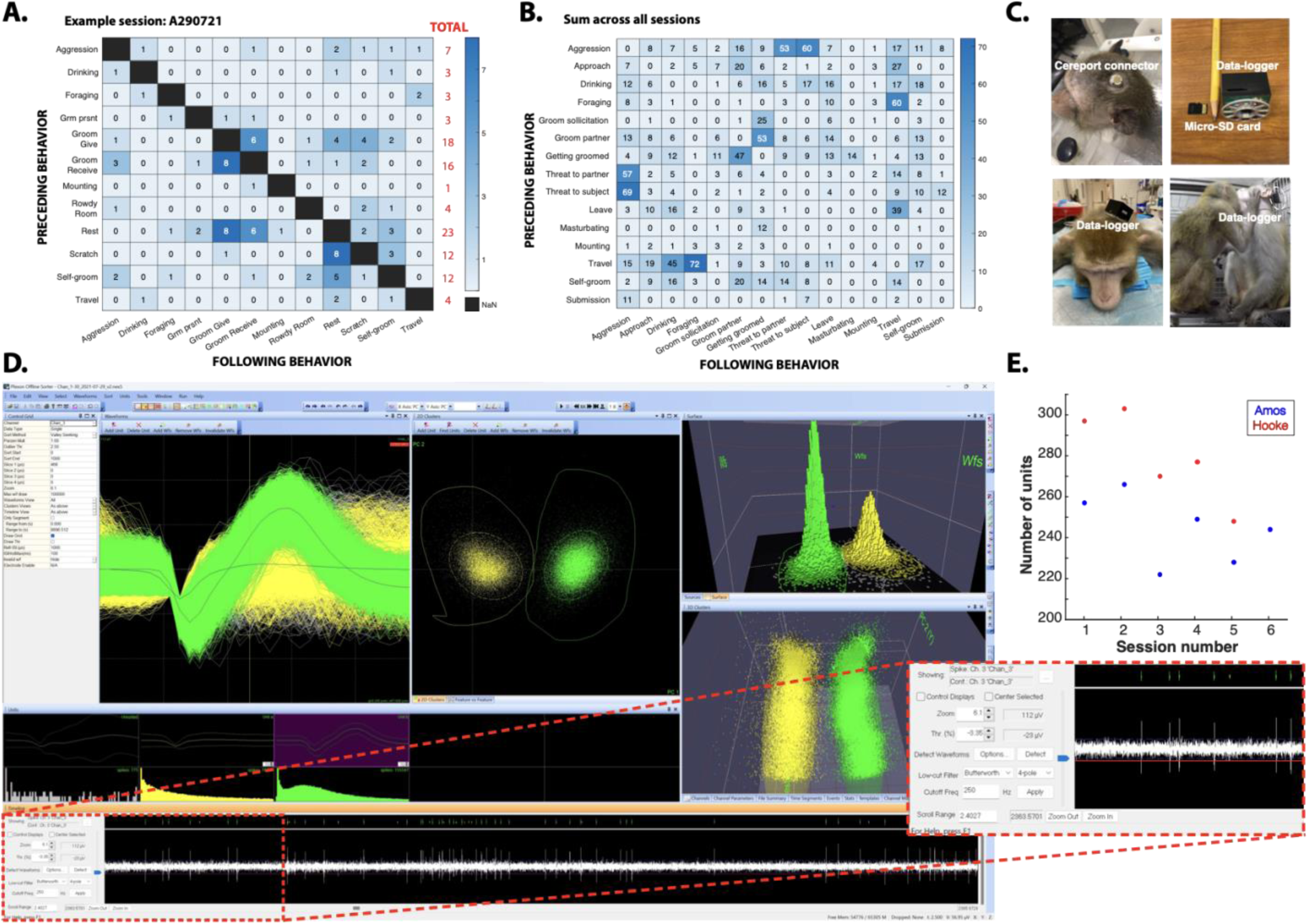
Behavior transition matrix, neurotechnology and neural data quality. (A) Absolute number of transitions from one behavior to the next for one example session. (B) Same as A) but across all sessions, monkeys pooled. (C) Images of wireless neural recording device. Note the minimal footprint of the cranial implant on the monkey’s head. (D) Example medium-amplitude unit (green) with average peak at 50uV. Bottom timeline plot (zoomed in) shows raw signal showing the high signal-to-noise ratio of spikes recording with the wireless logger. Red line indicates the threshold used, which is 3.7SD from noise amplitude. Only spikes crossing that line compose both the yellow and green clusters above. Noise typically has a low amplitude of +/- 15uV. Bottom right quadrant shows principal components vs time, used to confirm stability of the waveform shape over the entire session. (E) Number of units recorded across vlPFC and TEO in our two subject monkeys in each session, chronologically. All sessions were recording within 2 months of implantation.

**Fig S2.**
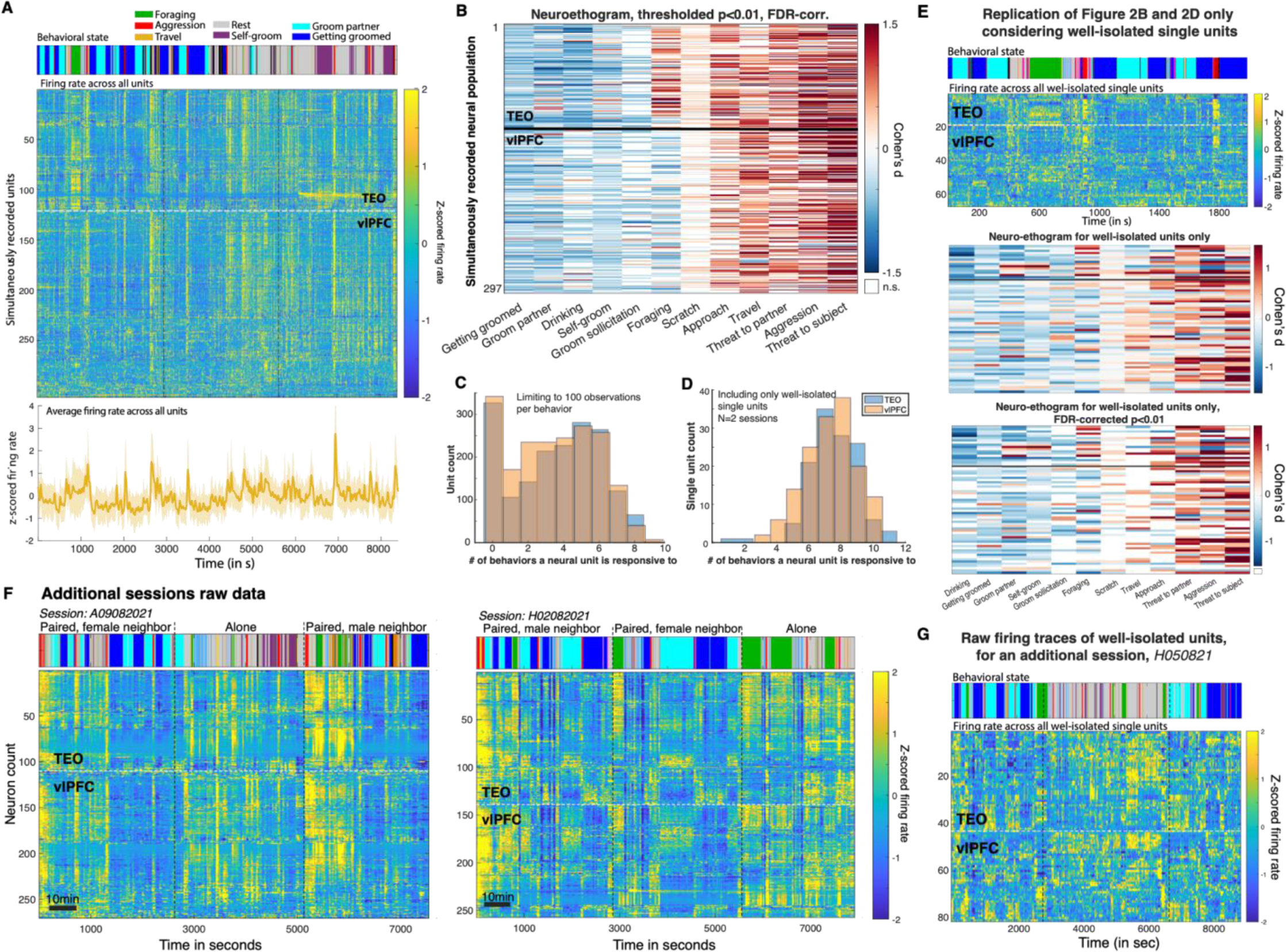
Single unit responses during behavior. (A) Top: behavior during example session; Middle: Z-scored firing rate of all units recorded in the same example session as Fig 2, ordered by activity pattern and units ordered using Rastermap ^37^; Bottom: Average z-scored firing across all recorded units in the same session. Shaded area is the standard deviation. (B) Neuro-ethogram from Fig 2D statistically thresholded at FDR-corrected *P* < 0.01. (C) Number of behaviors a neuron is selective for if limiting to 100 observations per behavior (sub-sampling 100 times through the data). (D) Same as Fig 2F but for well-isolated single units only for two sessions (one in each monkey). (E) Same as Fig 2B-D & S2B but only considering fully-isolated single units, as defined in Methods. (F) Raw firing aligned to behavior for two additional sessions to the one in Fig 2B and S2A, one from each monkey. Each row is an individual neural unit, ordered using the Rastermap toolbox^37^. Firing rates are Z-scored to maximum firing rate for each unit. Color bar on top represents behaviors performed over time with the same color code as in Fig 2B. (G) Z-scored raw firing aligned to behavior for only fully-isolated units for another session than in E, from the other monkey.

**Fig S3.**
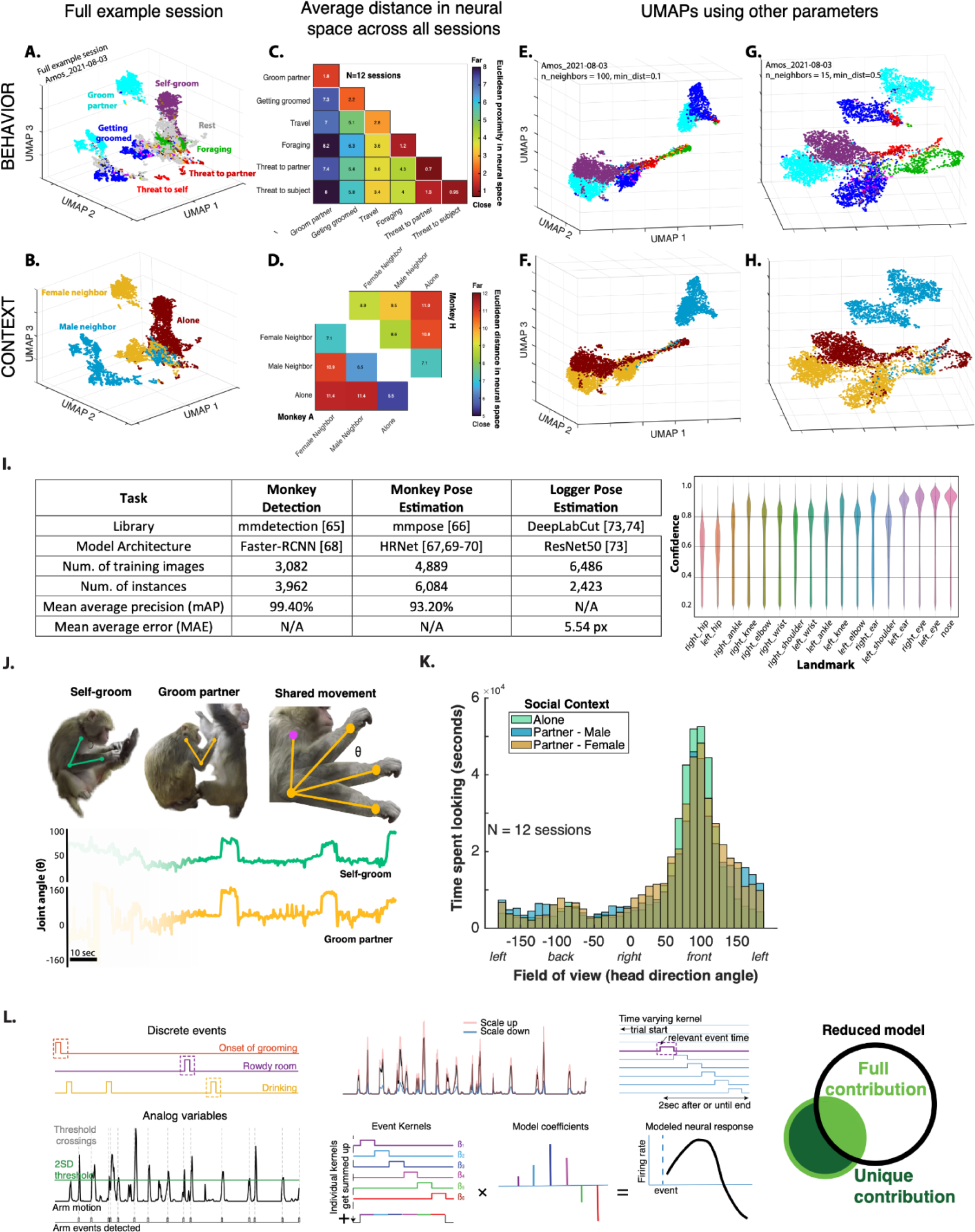
Neural population states segregate by behavior and context across sessions and are not reducible to visuo-motor contingencies. (**A-B**) Whole session UMAP visualization (same session as in **Fig 2A-D**). (**C)** Average distance between neural states across behaviors in UMAP space. Pooled across sessions and monkeys. (**D**) Average distance between neural states across social contexts. (**E-H**) UMAP plots as in Fig 2A but with different parameter values. Results remain qualitatively the same. (**I**) Left panel: Table summarizing behavior quantification models and their accuracies for monkey detection, monkey pose estimation, and datalogger pose estimation. Right panel: Confidence score by body landmark thresholding at c = 0.2. (**J**) Comparison of elbow joint angle for two behaviors, self-groom and groom partner. Top: depiction of the stereotypic self-groom behavior (left), groom partner (middle), and the shared motion pattern of shoulder-elbow-wrist across both behaviors (right). Bottom: plot of joint angle *θ* over time for self-grooming (green) and groom partner (yellow). Shaded portion indicates onset and offset of both behaviors. (**K**) Histograms of field of view across social contexts: alone, paired female neighbor, paired male neighbor. Field of view varied substantially within a social context and overlapped across contexts, decorrelating social context coding from visual inputs. (**L**) Schematic of ridge regression model to test the relative importance of behavior vs. movements and field of view to explain neural variance (**Fig 3I**).

**Fig S4.**
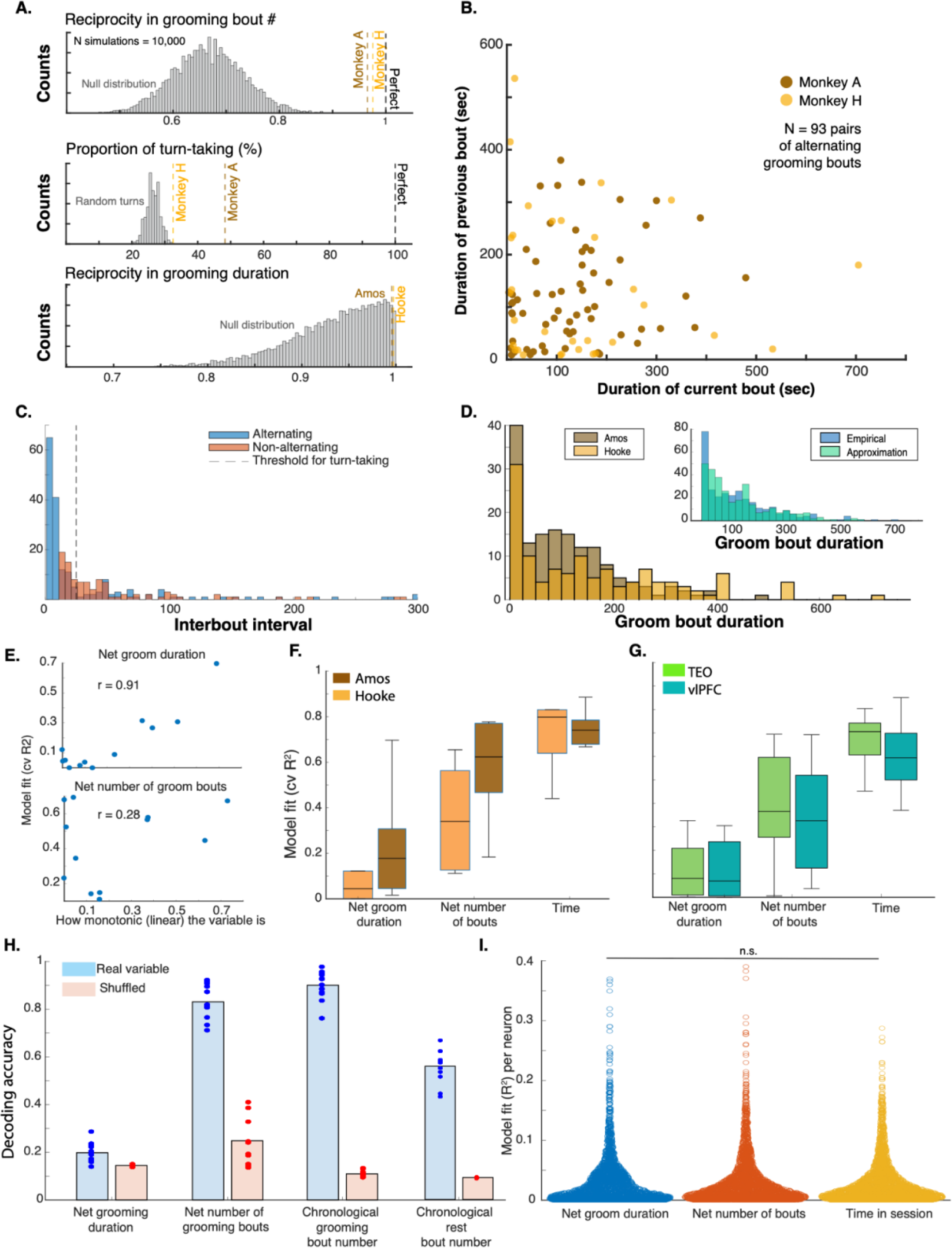
Behavioral evidence for grooming reciprocity and underlying neural correlates. (**A**) Simulations assuming monkeys were grooming randomly. 1 = perfectly reciprocal, 0 = fully one-sided. Simulations were run 10,000 times to obtain likelihood of the observed reciprocity if monkeys were not intentionally trying to reciprocate number of bouts (top), engage in turn-taking (middle) or reciprocate grooming duration (bottom) (see **Methods**). (**B**) Correlation in duration of consecutive alternating grooming bouts (e.g. I groom you, then you groom me). (**C**) Interbout interval for alternating grooming bouts (Give-Receive-Give, blue) vs non-alternating (Give-Give or Receive-Receive, red). Alternating bouts showed a bi-modal distribution with short inter-bout intervals <20s and longer ones. Based on this distribution we considered alternating bouts spaced by less than 20s to be “Turn-taking”. (**D**) Distribution of grooming bout duration for Amos (brown) vs. Hooke (yellow). Inset: Empirical distribution of grooming bout durations (blue) and approximated distribution (turquoise). Approximation uses an exponential distribution with empirical mean bout length as a parameter. (**E**) Correlation between ridge regression model fit and how linear & monotonic the response variable is. Top: Net groom duration; Bottom: Net number of grooming bouts. (**F**) Ridge regression model fit (cross-validated R^2^) for net grooming duration, net number of bouts and time combining separated by monkey (brain areas pooled). (**G**) Ridge regression model fit (cross-validated R^2^) for net grooming duration, net number of bouts and time combining separated by brain areas (monkeys pooled). (**H**) Cross-validation (5-fold) linear decoding accuracy of (from left to right): categorical net groom duration, categorical net number of bouts, chronological grooming bout number, chronological rest bout number. Red: shuffled control. (**I**) Distribution of single neuron cross-validated ridge regression fit for net groom duration, net groom bouts and time. Distributions are not statistically distinguishable.

**Fig S5.**
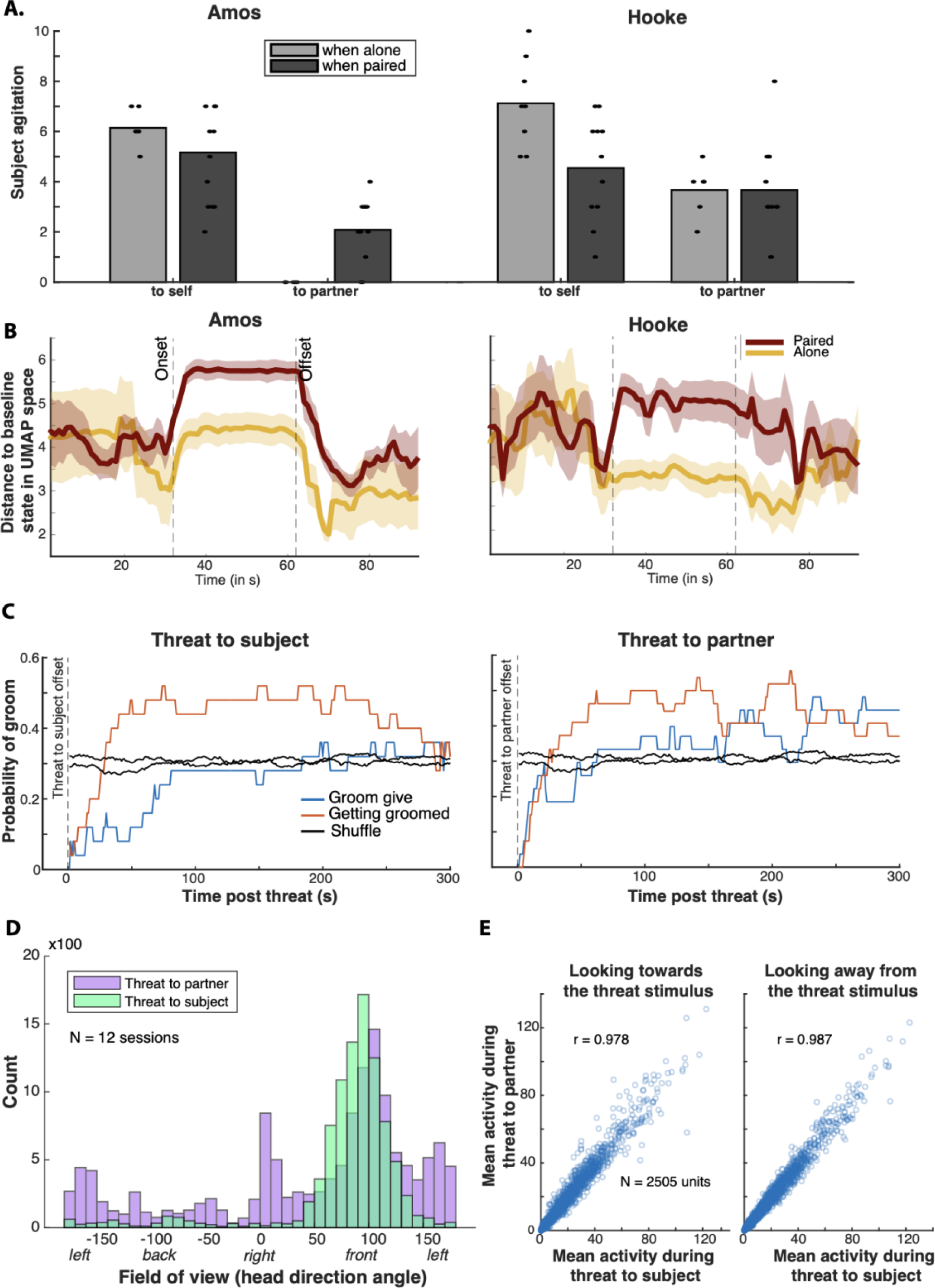
Neural signatures of social support and empathy in response to human threats. (**A**) Agitation score in response to human threats as in Fig 5B separated by monkey. (**B**) Same as in Fig 4C separated by monkey. (**C**) Probability of grooming given or received in the 5min following a threat. (**D**) Head direction during threats to self vs. partner, when paired. (**E**) Correlation in average neural responses to threats towards the subject vs. his partner when the subject is looking in front (left), towards the threatening stimulus, or not (right). Each point represents one unit, all units across sessions with video data were included (10 sessions). Note that, because of missing head direction data during threat bouts, we averaged neural activity across all threat data points and lost temporal variation occurring within a bout. This explains why we obtained an even higher correlation than calculated in Fig 5D.

**Table S1.**
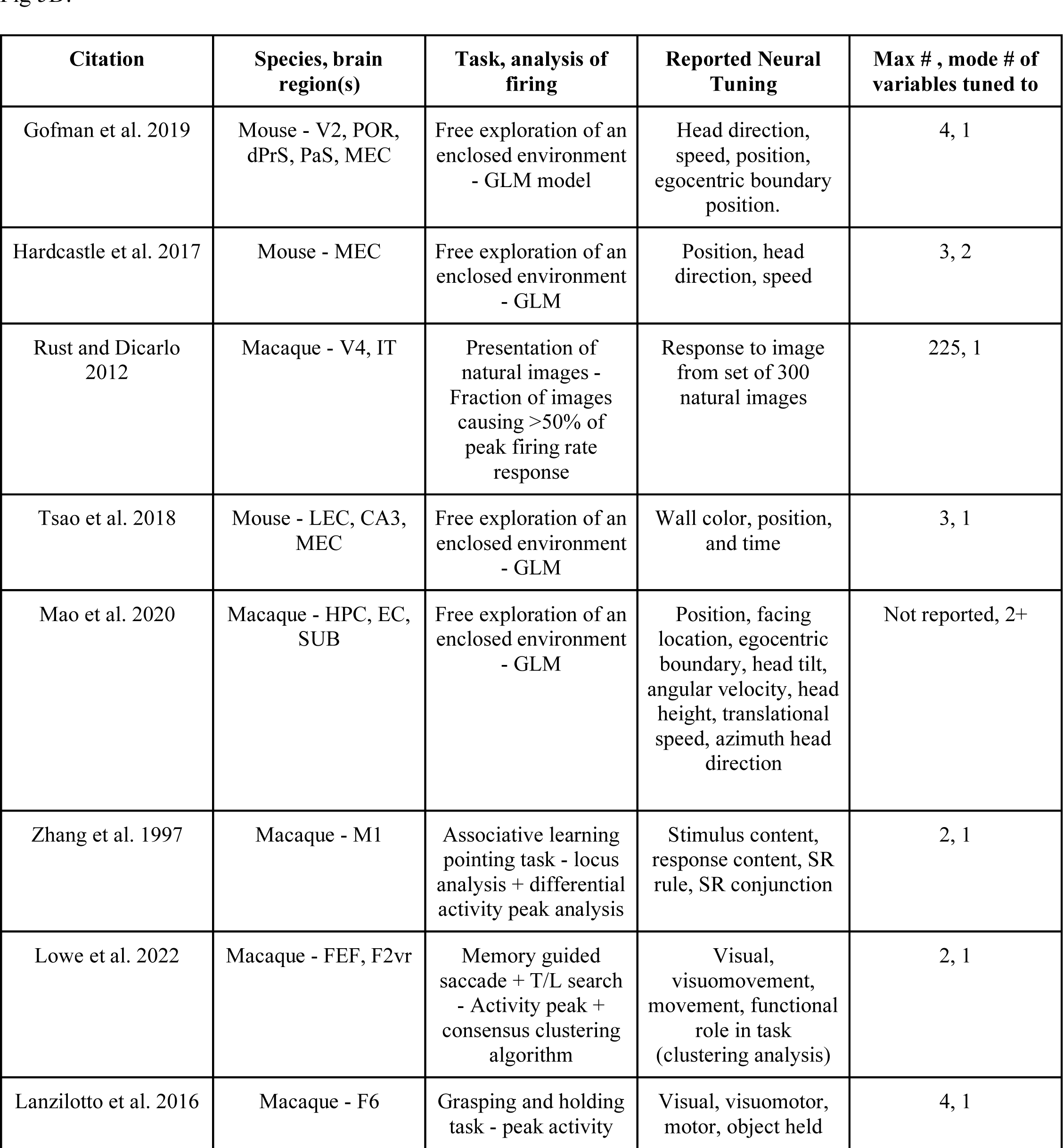
Selective literature review on sparse coding from neurophysiological data collected in traditional laboratory settings.

| Citation | Species, brain region(s) | Task, analysis of firing | Reported Neural Tuning | Max # , mode # of variables tuned to |
| --- | --- | --- | --- | --- |
| Gofman et al. 2019 | Mouse - V2, POR, dPrS, PaS, MEC | Free exploration of an enclosed environment<br>- GLM model | Head direction, speed, position, egocentric boundary position. | 4, 1 |
| Hardcastle et al. 2017 | Mouse - MEC | Free exploration of an enclosed environment<br>- GLM | Position, head direction, speed | 3, 2 |
| Rust and Dicarlo 2012 | Macaque - V4, IT | Presentation of natural images - Fraction of images causing >50% of peak firing rate response | Response to image from set of 300 natural images | 225, 1 |
| Tsao et al. 2018 | Mouse - LEC, CA3, MEC | Free exploration of an enclosed environment<br>- GLM | Wall color, position, and time | 3, 1 |
| Mao et al. 2020 | Macaque - HPC, EC, SUB | Free exploration of an enclosed environment<br>- GLM | Position, facing location, egocentric boundary, head tilt, angular velocity, head height, translational speed, azimuth head direction | Not reported, 2+ |
| Zhang et al. 1997 | Macaque - M1 | Associative learning pointing task - locus analysis + differential activity peak analysis | Stimulus content, response content, SR rule, SR conjunction | 2, 1 |
| Lowe et al. 2022 | Macaque - FEF, F2vr | Memory guided saccade + T/L search<br>- Activity peak + consensus clustering algorithm | Visual, visuomovement, movement, functional role in task (clustering analysis) | 2, 1 |
| Lanzilotto et al. 2016 | Macaque - F6 | Grasping and holding task - peak activity | Visual, visuomotor, motor, object held | 4, 1 |

